# Single-site DNA methylation activates retron defence by enabling effector tetramerisation

**DOI:** 10.64898/2026.10.13.758329

**Authors:** Panagiotis Spatharas, Alessio Ling Jie Yang, Edukondalu Mullapudi, Grzegorz Chojnowski, Federico Corona, Devansh Kumar, Jacob Bobonis, Athanasios Typas, Matthias Wilmanns

## Abstract

Retrons are unique among bacterial defence systems in using DNA to sense phage infection. They achieve this by producing multicopy single-stranded DNA (msDNA) that monitors the activity of DNA-targeting phage proteins. Although we understand how extensive msDNA perturbations such as degradation activate retrons, activation by subtle chemical modifications remains enigmatic. Here, we used a structure-function approach to elucidate how single-site msDNA methylation activates Retron-Sen2. The cryo-EM structure of Retron-Sen2 features RcaT, the most prevalent retron effector, sequestered as an inactive dimer within a four-component assembly of previously undescribed architecture. Methylation by DNA adenine methyltransferase disrupts the RcaT-msDNA interaction and exposes a previously occluded interface that enables RcaT dimers to form tetramers, leading to retron disassembly. We show that RcaT dimer–dimer assembly is essential for its toxicity. To assess whether this is a conserved RcaT feature, we characterised additional homologues. RcaTs capable of tetramerisation or higher-order oligomerisation are toxic and form distinct phylogenetic clades from RcaTs that remain dimeric and are non-toxic. Together, our findings provide a paradigm for retron activation by single-site msDNA modifications and showcase how variation in an intrinsic protein property can diversify the toxicity of the same bacterial defence effector during evolution.

## Introduction

Retrons were the first retroelements discovered in bacteria^1,2^. They encode a specialised reverse transcriptase (RT) that synthesises multicopy single-stranded DNA (msDNA), a branched DNA/RNA hybrid whose DNA and RNA components are termed msd and msr, respectively^3^. Only recently was it established that the physiological role of retrons is to defend against phage infection^4–6^. This is enabled by diverse effector proteins^4–6^ encoded adjacently to the RT and msDNA. Most characterised retrons share a common organisational principle: the RT binds msDNA, forming a complex that neutralises the effector in the absence of infection^7–13^. What sets retrons apart from other bacterial defence systems is their use of msDNA as an infection sensor to detect the activity of phage-encoded proteins on DNA^6^. This triggers their effector proteins to promote abortive infection^4–6^ via mechanisms that remain only partially understood.

Retrons are classified into at least 13 distinct types based on their effector diversity^14^. Recent studies have begun to reveal that this diversity extends to the structures and activation mechanisms of retron assemblies^7–13^. Two such mechanisms have been characterised: msDNA-mediated recognition of phage single-stranded DNA-binding proteins (SSBs)^11,15,16^ and degradation of msDNA by phage nucleases^7,9,17^. These mechanisms involve extensive perturbations of the msDNA, which is an integral part of retron assemblies^6^. Retron activation can also be triggered by much subtler modifications, such as single-site msDNA methylation by phage-encoded methyltransferases^6,18,19^. Yet how such a minimal perturbation alters large retron assemblies to activate phage defence remains obscure.

Type II retrons constitute the most abundant retron family^5^. Several members, including Sen2, Eco9 and Eco1, are triggered by phage-encoded methyltransferases^6,18,19^. They are characterised by effectors that share a conserved architecture, comprising a nucleoside deoxyribosyltransferase-like (NDT-like) domain and a DNA-binding domain^5,6,14^. Type II effectors are termed retron cold-anaerobic toxins (RcaTs), after the conditional activity of the first characterised member, Sen2^6^. Some of these effectors function as bona fide toxins (e.g. Sen2, Eco9, Eco10), with the RT-msDNA complex serving as the antitoxin of a toxin–antitoxin system^6^. Other RcaTs are non-toxic (Eco1, Eco3) despite conferring effective phage defence^6^. In Retron-Eco1, this defence is linked to NAD⁺ hydrolysis and auto-ADP-ribosylation^13,18^. The molecular basis for the differential toxicity of RcaT effectors is unclear.

Here, we use Retron-Sen2^20^ (Fig. 1a) as a model system to elucidate how msDNA methylation activates retron immunity and investigate the basis of differential RcaT toxicity. We first determine the structure of Retron-Sen2, in which RcaT is sequestered as an inactive dimer within a complex that can further assemble into filaments. We then show that methylation at a single GATC motif in the msDNA by phage DNA adenine methyltransferases (Dam) promotes complex disassembly and enables RcaT to transition from the inactive dimer to toxic higher-order oligomers. This transition is essential for RcaT toxicity and is prevented by the retron antitoxin through interactions with msDNA. We finally show that higher-order oligomerisation beyond the intrinsic dimer is a hallmark of toxic RcaTs, whereas non-toxic homologues are dimeric across the RcaT phylogeny.

**Figure 1.**
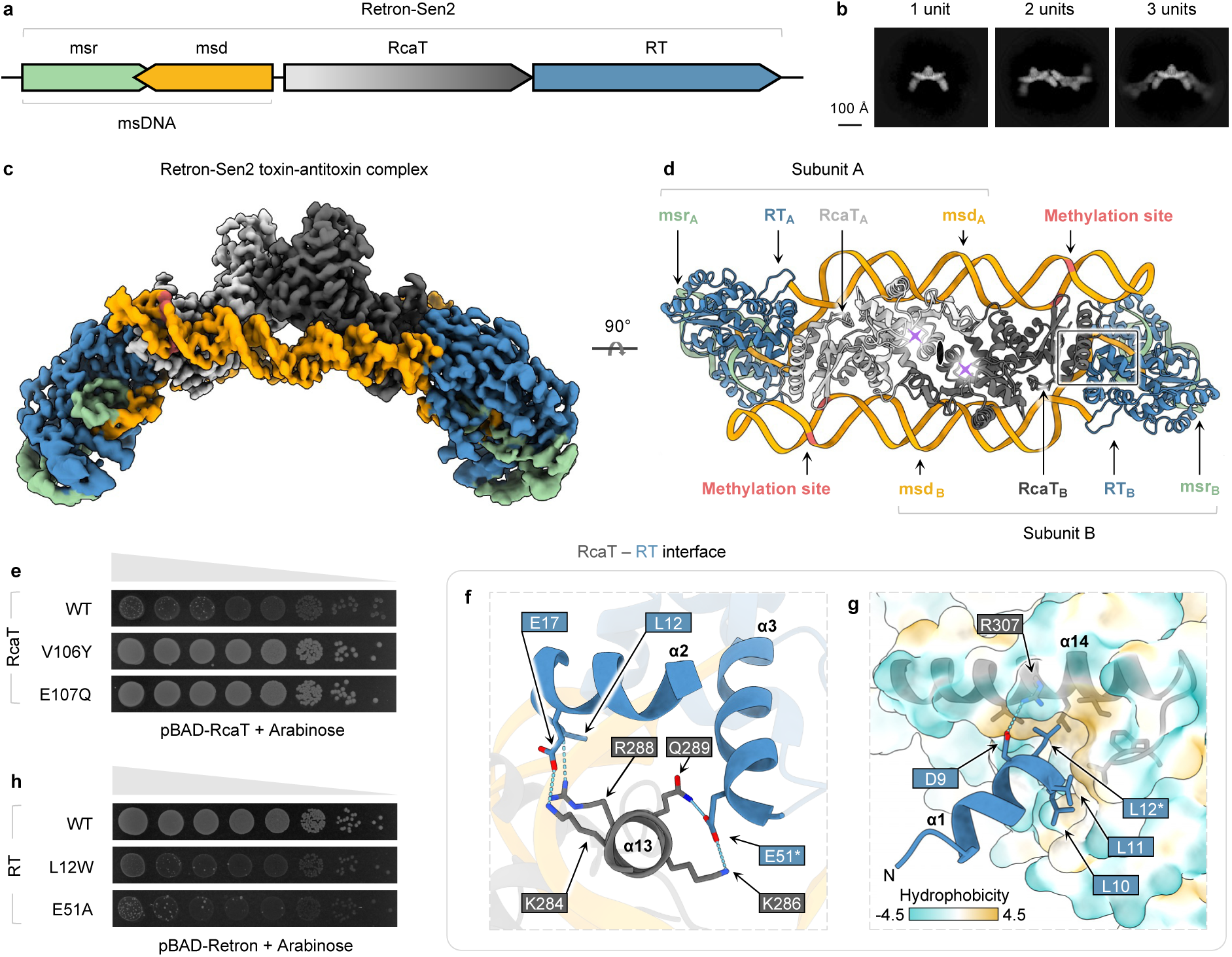
The Retron-Sen2 toxin–antitoxin complex sequesters RcaT dimers. **a,** The Retron-Sen2 locus encodes RcaT (grey), a reverse transcriptase (RT; blue) and a non-coding RNA comprising msr (green) and msd (orange). The RT reverse-transcribes the msd region to produce msDNA, a DNA–RNA hybrid comprising msd (DNA) and msr (RNA). The RT and the msDNA form an antitoxin module that inhibits RcaT6. The colour code applies throughout unless otherwise indicated. **b**, Cryo-EM 2D class averages show a single toxin–antitoxin complex unit (left) and filaments containing two (middle) or three (right) units. Scale bar, 100 Å. **c**, Cryo-EM map of the toxin–antitoxin complex at 3.37 Å resolution. **d**, Atomic model of the toxin–antitoxin complex rotated by 90° relative to panel c shows two identical subunits (A, B) related by twofold symmetry (central ellipse). Each subunit contains RcaT, RT, msr and msd in a 1:1:1:1 ratio. The central RcaT dimer (monomers as light and dark grey) is encased by two antitoxin modules (RT-msd-msr). Each msd contacts RcaT within its own subunit (proximal RcaT) and the opposite subunit (distal RcaT). Lilac asterisks mark RcaT active sites; red marks msDNA methylation sites. The box highlights the RcaT–RT interface detailed in f and g. **e**, Mutations disrupting RcaT dimerisation (V106Y) or catalytic activity (E107Q) abolish toxicity, as shown by tenfold serial dilution spot assays of strains expressing RcaT from a pBAD plasmid with arabinose induction. **f**, Close-up of the RcaT–RT interface. Dashed lines indicate putative hydrogen bonds, and asterisks mark residues selected for mutagenesis throughout. **g**, The RcaT–RT interface also involves a hydrophobic pocket formed by RcaT helix α14 that accommodates RT residues L10, L11, and L12, and an electrostatic interaction between RcaT R307 and RT D9. The RcaT surface is coloured according to the Kyte–Doolittle hydrophobicity scale. **h**, Mutations in the RcaT–RT interface (RT L12W and RT E51A) disrupt inhibition of RcaT by the antitoxin and induce toxicity in strains expressing Retron-Sen2 (n = 3 biological replicates) – spot assays as in panel e.

## Results

### The Retron-Sen2 toxin–antitoxin complex sequesters RcaT dimers

To reconstitute the Retron-Sen2 toxin–antitoxin complex, we tagged the RT and used it to co-purify msDNA together with a catalytically inactive RcaT-E107Q variant to prevent leaky toxicity^6^ (Extended Data Fig. 1a). We then determined the structure of the resulting complex by single-particle cryo-EM (Extended Data Fig. 1b-c). The analysis revealed that Retron-Sen2 forms a core toxin–antitoxin complex that can further assemble into filaments (Fig. 1b).

The core toxin–antitoxin complex was resolved to a global resolution of 3.37 Å (Fig. 1c). It is composed of two equivalent 1:1:1:1 RcaT-RT-msd-msr subunits (A, B), which are related by twofold symmetry (Fig. 1d). The two subunits assemble through a central RcaT dimer, encased by two flanking antitoxin modules (RT-msd-msr). The msd not only contacts RcaT from its own subunit (proximal), but also the RcaT from the opposite subunit (distal). The msr is truncated and interacts exclusively with the RT and msd from the same subunit (Extended Data Fig. 2). The presence of this msr fragment was unexpected, as Retron-Sen2 msDNA was thought to be RNA-free, since ExoVII-mediated debranching is required for its maturation^6,21^.

RcaT dimerisation is mediated by the NDT-like domain (Extended Data Fig. 3a), which harbours the catalytic triad^13^—F28, E85, and E107 (Extended Data Fig. 3b). The dimerisation interface is dominated by an extended hydrophobic core (Extended Data Fig. 3c), which is a common feature of NDT dimers^22^. Both the catalytic triad and most residues contributing to dimerisation are conserved across the RcaT phylogeny (Extended Data Fig. 3d). Disrupting the catalytic core (E107Q) or the dimerisation interface (V106Y) abolished RcaT toxicity (Fig. 1e), underscoring their functional importance.

RcaT is kept inactive through interactions with the RT and msd. Its DNA-binding domain faces outward from the dimer and engages the N-terminus of the RT (Extended Data Fig. 3a), burying approximately 600 Å² of surface area on each side (Supplementary Data 1). RcaT helix α13 interacts with RT helices α2 and α3, including hydrogen bonds between RcaT residues K286 and Q289 and RT residue E51 (Fig. 1f). RcaT helix α14 and its preceding loop form a hydrophobic pocket that accommodates RT residues L10–L11–L12 and features an electrostatic interaction between RcaT R307 and RT D9 (Fig. 1g). Mutations at the RcaT–RT interface (RT L12W and RT E51A) disrupt RcaT inhibition and induce toxicity (Fig. 1h), highlighting the essential role of RT in RcaT neutralisation.

The DNA-binding domain of RcaT also engages the msd stem–loop (Fig. 2a). Multiple RcaT–msd contacts are formed on each side of the dimer, together burying approximately 2,600 Å² of surface area (Supplementary Data 1). Contacts with the proximal RcaT are localised primarily near the base of the msd stem. In contrast, the distal RcaT contributes interactions clustered closer to the msd loop, with most localising to the region containing the GATC methylation motif.

**Figure 2.**
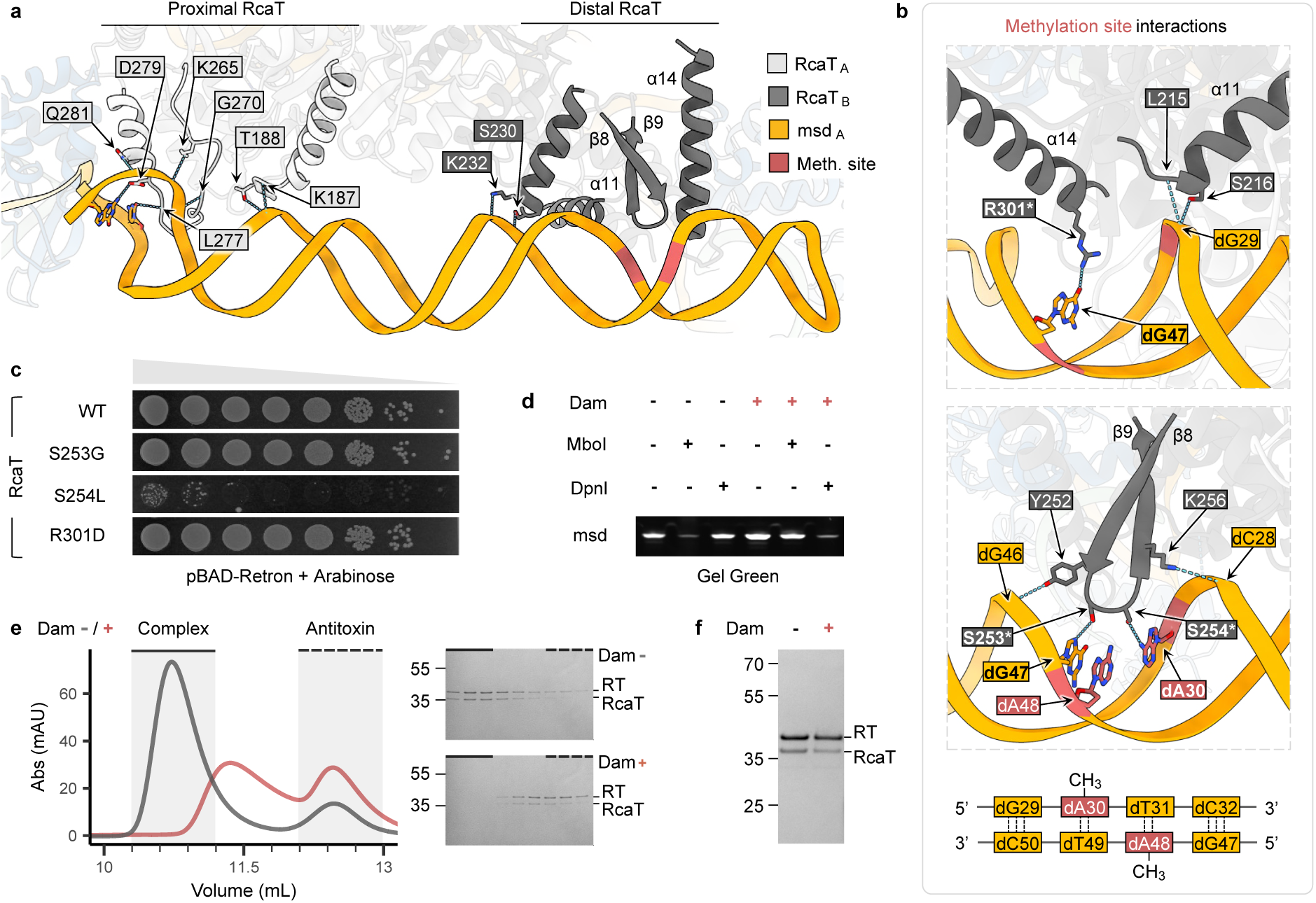
Methylation of msDNA promotes Retron-Sen2 toxin–antitoxin complex disassembly and RcaT release. **a**, Proximal (light grey) and distal (dark grey) RcaT engage msd (orange) asymmetrically. **b**, Distal RcaT interacts with the msd methylation site (red). RcaT helices α11 and α14 (upper panel) and the β8–β9 hairpin (lower panel) contact the Dam-recognised GATC motif, including the methylatable adenines dA30 and dA48 (bottom schematic). **c**, Of three substitutions in RcaT residues contacting the methylation site, S254L abolishes inhibition of RcaT by the antitoxin and induces toxicity in strains expressing Retron-Sen2, whereas S253G and R301D do not (n = 3 biological replicates) – spot assays as in Fig. 1. **d**, Digestion assays with methylation-sensitive enzymes demonstrate msDNA methylation after *in vitro* Dam treatment, as shown by Gel Green-stained urea–PAGE. MboI cleaves unmethylated GATC sites, whereas DpnI cleaves methylated sites. **e**, SEC profiles at 280 nm compare methylated (red) and unmethylated (grey) Retron-Sen2 samples (representative of n = 2 biological replicates). Methylation eliminates the toxin–antitoxin complex peak and produces a new peak at a later elution volume, accompanied by an increase in the antitoxin peak. The new peak contains both RT and RcaT, as shown by SDS–PAGE of the fractions. Solid and dotted lines indicate fractions corresponding to the toxin–antitoxin complex and free antitoxin, respectively, on both the SEC profiles and SDS–PAGE gels. **f**, msDNA methylation reduces RcaT pull-down by Strep-tagged RT, indicating partial release of RcaT from the toxin–antitoxin complex (representative of n = 2 biological replicates).

We next examined the filament assemblies observed in our initial cryo-EM analysis (Extended Data Fig. 1b). The toxin–antitoxin complex serves as the repeating unit of these filaments (Fig. 1b). For further structural analysis, we used a map of a two-unit filament (Extended Data Fig. 4a). Adjacent toxin– antitoxin complexes are connected through a filament interface formed by an RT-msDNA antitoxin dimer. The truncated msr appears to contribute to this dimer interface (Extended Data Fig. 4b-c). Filament formation has also been reported for the homologous Retron-Eco1^13,18^, but its architecture differs markedly from that observed here (Extended Data Fig. 4d-e). In Retron-Eco1, the core complex features an RT-msDNA dimer bound to two independent RcaT monomers, whereas the filament interface is formed by an RcaT dimer. This arrangement is reversed in Retron-Sen2, where RcaT dimerisation occurs within the core complex and RT-msDNA dimerisation forms the filament interface. Surprisingly, the Retron-Sen2 RT-msDNA antitoxin is monomeric without RcaT (Extended Data Fig. 5), indicating that antitoxin dimerisation is specific to the filament context.

### RcaT interacts with the methylation site of msDNA

We then used the Retron-Sen2 toxin–antitoxin complex structure to understand how site-specific adenine methylation by Dam triggers activation^6^. The GATC methylation motif of msd spans dG29– dC32 and the complementary bases dG47–dC50 (Fig. 2b, bottom schematic). Dam recognises this motif and methylates dA30 and dA48, thereby activating the system^6^. RcaT engages the methylation motif through helices α11 and α14 and the β8–β9 hairpin (Fig. 2a). In helix α11, the backbone amide of L215 and the side chain of S216 interact with the phosphate group of dG29 (Fig. 2b, upper panel). Helix α14 contributes R301, whose side chain forms a hydrogen bond with the base of dG47 (Fig. 2b, upper panel), conferring sequence-specific recognition. The β8–β9 hairpin inserts directly into the major groove of the methylation motif (Fig. 2b, lower panel). Specifically, Y252 and K256 engage the phosphate groups of dG46 and dC28, which flank the motif, thereby locking residues S253 and S254 directly within the major groove. S253 is positioned to form a hydrogen bond with the base of dG47, contributing further sequence specificity. S254 lies directly between the two methylatable adenines, dA30 and dA48, and is well positioned to hydrogen bond with dA30 (Fig. 2b, lower panel). Surprisingly, S253G and R301D substitutions failed to induce toxicity (Fig. 2c), suggesting that these residues may serve additional roles beyond msDNA binding (see below for their role in RcaT dimer–dimer assembly). In contrast, introducing the bulky S254L substitution to mimic the steric effect of dA30 methylation induced cellular toxicity (Fig. 2c). These observations support a model in which methylation-induced steric hindrance disrupts the RcaT–msd interface, ultimately leading to toxin activation.

### Methylation of msDNA releases RcaT from the toxin–antitoxin complex

To further test the effect of methylation on the RcaT–msd interface, we subjected the purified Retron-Sen2 toxin–antitoxin complex to *in vitro* methylation using *E. coli* Dam, which phenocopies the effect of phage P1 Dam^6^. Methylation of msDNA was confirmed using the methylation-sensitive restriction enzymes MboI and DpnI, which cleave only unmethylated or methylated GATC sites, respectively (Fig. 2d). Equal amounts of methylated and unmethylated complexes were then analysed by size-exclusion chromatography (SEC; Fig. 2e). The unmethylated sample eluted predominantly as intact toxin– antitoxin complex, with a smaller peak corresponding to free antitoxin. In contrast, methylation resulted in loss of the toxin–antitoxin complex peak and appearance of a new, later-eluting peak containing both RcaT and antitoxin, accompanied by a substantial increase in the free antitoxin peak. These changes demonstrate disassembly of the toxin–antitoxin complex, with antitoxin dissociation and the possibility of RcaT release. To directly assess RcaT release, methylated and unmethylated samples were subjected to a second round of affinity purification via the RT affinity tag (Fig. 2f). In the unmethylated sample, all components co-eluted, consistent with an intact toxin–antitoxin complex. Upon methylation, RcaT still co-eluted with the antitoxin but at reduced levels. Together, these results confirm that msDNA methylation promotes disassembly of the Retron-Sen2 toxin–antitoxin complex and allows for partial RcaT release.

### RcaT forms toxic tetramers

To understand what happens to released RcaT, we purified RcaT in the absence of the antitoxin, using the catalytically inactive RcaT-E107Q mutant. SEC yielded a profile consistent with a tetrameric species (Extended Data Fig. 6a). Blue native PAGE (BN-PAGE) further revealed a higher-molecular-weight species, indicative of higher-order oligomerisation (Extended Data Fig. 6a). To probe the reported temperature dependence of RcaT-Sen2 activity^6^, we subjected purified RcaT to nanoscale differential scanning fluorimetry (nanoDSF). The onset of aggregation was observed only above 52.5 °C (Extended Data Fig. 6b), indicating that the reduced *in vivo* toxicity previously reported at 37 °C is unlikely to arise from compromised protein stability.

We then determined the structure of RcaT by cryo-EM (Extended Data Fig. 6c-d). Consistent with the SEC data, the dominant species was a tetramer arranged as two stacked dimers, which we reconstructed to a global resolution of 3.19 Å (Fig. 3a-b). We also observed higher-order oligomers formed by the stacking of additional RcaT dimers. From these particles, we reconstructed an RcaT decamer and locally refined its three central dimers to obtain a higher-resolution RcaT hexamer (Extended Data Fig. 6e). In all oligomeric states, dimer stacking was mediated by the same dimer– dimer interface (Fig. 3c). For subsequent structural and functional analysis, we used the predominant tetrameric assembly as the reference structure.

**Figure 3.**
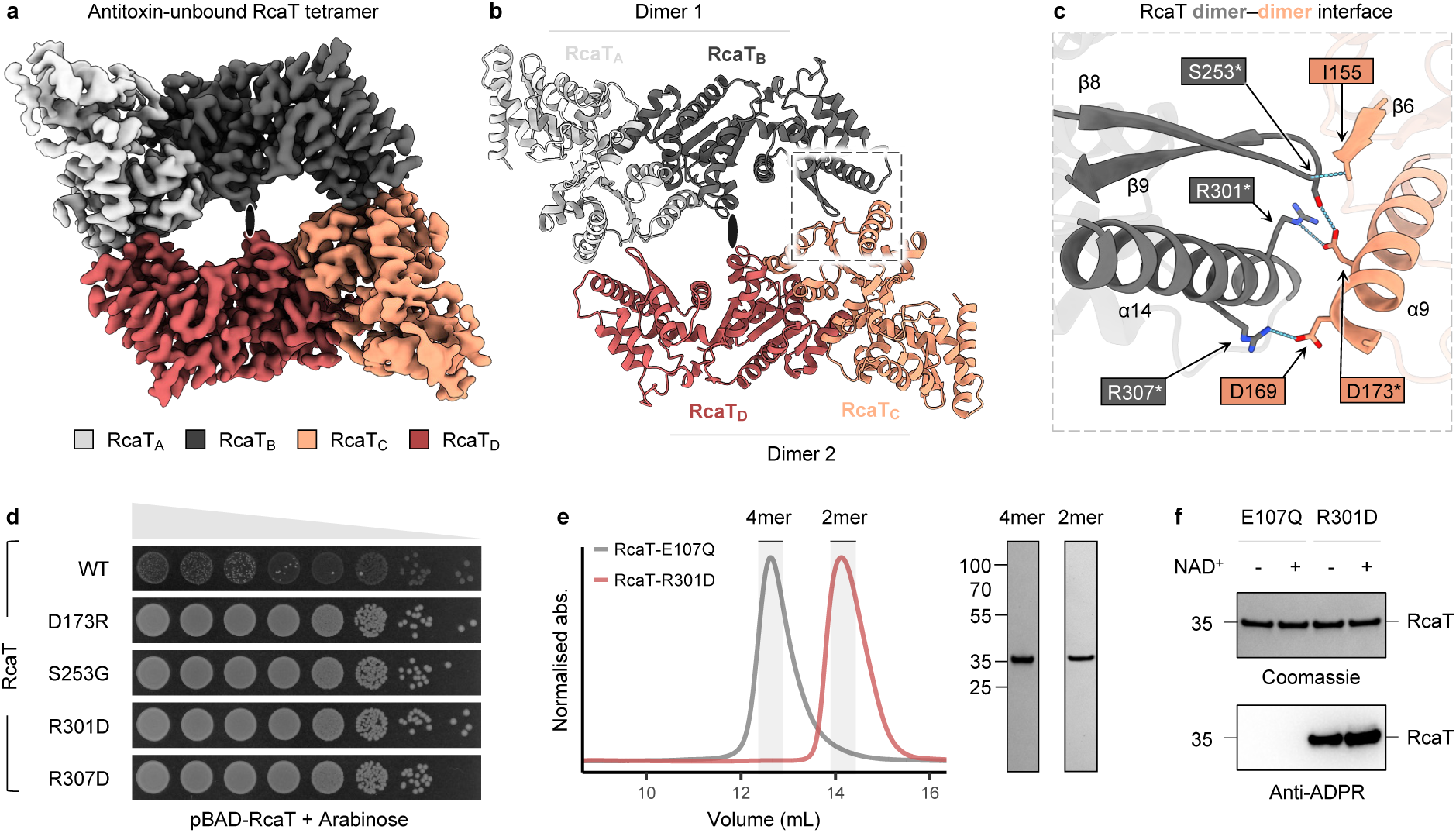
RcaT forms toxic tetramers. **a,** A 3.19 Å cryo-EM map and **b**, atomic model reveal that antitoxin-free RcaT forms a tetramer arranged as a symmetric dimer of dimers in light/dark grey and light/dark salmon. The central ellipse marks the position of the twofold symmetry axis. Each dimeric module resembles the RcaT dimer observed in the toxin–antitoxin complex (Fig. 1d). The box highlights the dimer–dimer interface detailed in c. **c**, The dimer–dimer interface involves the same secondary-structure elements and residues that mediate RcaT interactions with the msd methylation site and RT in the toxin– antitoxin complex (Fig. 2b and Fig. 1g, respectively). **d**, Dimer–dimer assembly is essential for toxicity. All tested dimer–dimer interface mutants (D173R, S253G, R301D and R307D) abolish toxicity (n = 3 biological replicates) – spot assays as in Fig. 1. **e**, The R301D substitution disrupts RcaT dimer–dimer assembly, yielding only dimers, as shown by SEC and SDS–PAGE of the corresponding fractions. **f**, The RcaT-R301D dimer retains auto-ADP-ribosylation activity despite loss of toxicity, as detected by anti-ADPR immunoblotting. Signal is detectable from endogenous levels of NAD+, but intensity increases upon addition of 1 mM NAD⁺. The catalytically inactive control RcaT-E107Q shows no detectable signal for auto-ADP-ribosylation activity.

The RcaT dimer–dimer interface is formed by the β8–β9 hairpin and helix α14 from one RcaT dimer engaging helix α9 from the opposing dimer (Fig. 3c). At the residue level, this interface is formed by polar interactions involving the side-chain pairs S253–D173, R301–D173, and R307–D169, as well as a backbone interaction between S253 and I155. Notably, the structural elements and residues mediating RcaT dimer–dimer assembly overlap extensively with those involved in binding the msd methylation site and, to a lesser extent, the RT in the toxin–antitoxin complex (Fig. 3c, Fig. 2b, Fig. 1g). This indicates that RcaT dimer–dimer assembly competes directly with antitoxin binding, rendering the two states mutually exclusive.

Introducing mutations (D173R, S253G, R301D, and R307D) within the dimer–dimer interface consistently abolished RcaT toxicity (Fig. 3d). This also explains why the S253G and R301D substitutions failed to induce toxicity despite perturbing RcaT interactions with the msd methylation site in the toxin–antitoxin complex (Fig. 2c). Although these substitutions may promote RcaT release, they simultaneously disrupt RcaT dimer–dimer assembly. To test this, we purified the RcaT-R301D mutant and confirmed that it yields predominantly dimeric species by SEC (Fig. 3e). Together, these results demonstrate that the dimer–dimer assembly is essential for RcaT toxicity. To assess whether dimer– dimer assembly is required for catalytic activity, we performed western blotting with an anti-ADPR antibody and found that RcaT-R301D retained auto-ADP-ribosylation activity, even at endogenous NAD⁺ levels (Fig. 3f). Thus, catalytic competence alone is seemingly insufficient for toxicity, implying that dimer–dimer assembly is required for a downstream step, such as target recognition.

To further explore how oligomeric state relates to toxicity, we compared the structure of the RcaT dimer within the antitoxin-unbound tetramer with that in the toxin–antitoxin complex. In the toxin– antitoxin complex, the RcaT dimer exhibits twofold symmetry (Extended Data Fig. 7a). By contrast, the dimer derived from the tetramer has its subunits related by a rotation of 167°. This asymmetry originates from a region proximal to the active site and is accompanied by substantial rearrangement of the putative NAD⁺-binding site identified in Retron-Eco1^13,18^ (Extended Data Fig. 7b). Whether this symmetry breaking contributes to RcaT activity remains uncertain. On the other hand, the requirement for dimer–dimer assembly is clear. We therefore asked whether the ability to oligomerise beyond the dimer distinguishes toxic from non-toxic RcaT homologues.

### Oligomeric state determines toxicity across the RcaT phylogeny

To test the hypothesis that non-toxic RcaTs are intrinsically dimeric, we first used AlphaFold-Multimer^23,24^ to assess the oligomeric state of the non-toxic RcaT-Eco1. Tetramer prediction yielded two independent dimers with a low inter-chain predicted template modelling score (ipTM = 0.37; Extended Data Fig. 8a). By contrast, AlphaFold predicted an RcaT-Sen2 tetramer with a high ipTM score (0.82) that closely matched the experimentally determined structure (Extended Data Fig. 8b). To validate these predictions experimentally, we purified wild-type RcaT-Eco1 and analysed its oligomeric state by SEC, which confirmed exclusive dimer formation (Fig. 4a). These observations prompted us to ask whether the RcaT oligomeric state correlates with its toxicity across its phylogenetic tree.

**Figure 4.**
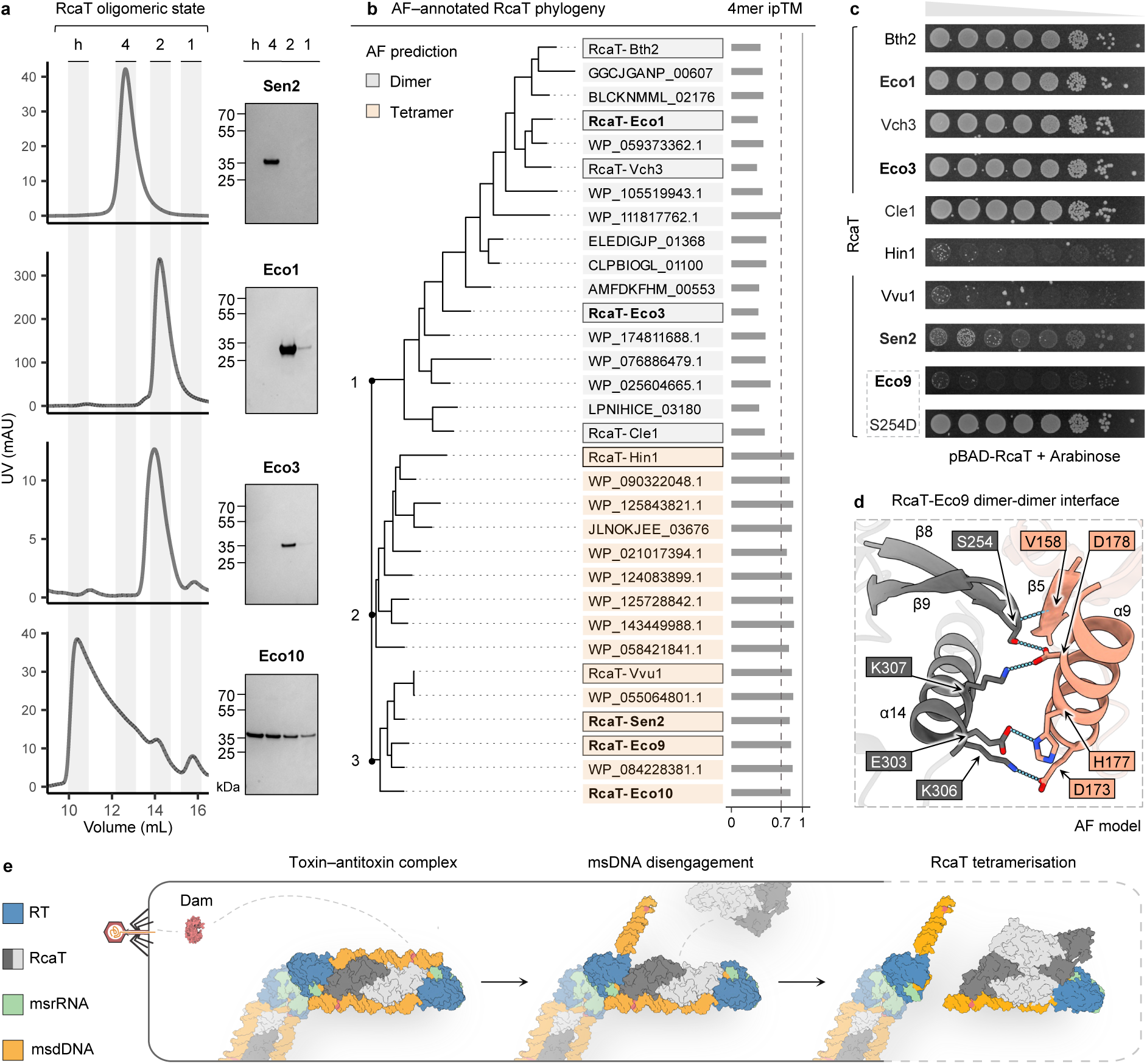
RcaTs diverged into toxic oligomeric and non-toxic dimeric variants over the course of evolution. **a,** Known RcaT homologues differ in oligomeric state, as determined by SEC. Shaded regions indicate higher-order oligomers (h), tetramers (4), dimers (2), and monomers (1). SDS–PAGE of the indicated fractions is shown at right. Toxic RcaT-Sen26 is predominantly tetrameric, non-toxic RcaT-Eco1 and RcaT-Eco36 are exclusively dimeric, and toxic RcaT-Eco106 forms species ranging from monomers to higher-order oligomers. **b**, RcaT phylogeny, annotated by AlphaFold predictions of oligomer assemblies, shows a clear divergence into dimer-(grey) and tetramer-forming (peach) RcaT clades. Bars indicate ipTM scores for predicted tetramers; models are confident (ipTM ≥ 0.7, dashed line) for all members of clades 2 and 3, but for none of clade 1. RcaTs tested for toxicity in panel c and those with known toxicity6 are outlined; the latter are also shown in bold. **c**, The toxicity of previously uncharacterised RcaTs can be inferred from AlphaFold predictions of their oligomeric state. Tetramer-predicted Vvu1 and Hin1 are toxic, whereas dimer-predicted Bth2, Vch3, and Cle1 are non-toxic. The phosphomimetic Eco9 S254D mutant loses toxicity (n = 3 biological replicates). Spot assays as in Fig. 1. **d**, The AlphaFold-predicted dimer–dimer interface of RcaT-Eco9 resembles that of RcaT-Sen2 (Fig. 3c). Eco9 S254 (mutated in panel c) aligns with Sen2 S253 at the interface. **e**, Proposed model: upon infection, phage-encoded Dam methylates the Retron-Sen2 toxin–antitoxin complex, triggering disengagement of msDNA from RcaT. The disengaged msDNA state is modelled after the alternative antitoxin conformation described in Extended Data Fig. 5. Loss of the RcaT–msDNA interface creates space for RcaT dimer–dimer assembly. Dimer– dimer assembly unleashes RcaT toxicity, targeting the infected host to curtail phage propagation.

We therefore selected 30 additional RcaTs spanning the previously described RcaT phylogeny^6^ (reproduced in Extended Data Fig. 8c) and constructed a representative subtree comprising three major clades (Fig. 4b). For each RcaT, we generated AlphaFold predictions of both dimeric and tetrameric assemblies (Supplementary Data 2-4). All RcaTs were predicted to form dimers with high confidence, indicating that dimerisation is conserved within the entire family (Extended Data Fig. 8d). In contrast, tetramer predictions revealed a clear division: while RcaTs in clades 2 and 3 showed consistently high tetramer ipTM scores, clade 1 RcaTs displayed uniformly low confidence for tetramer formation (Fig. 4b). Consistent with this division, non-toxic RcaT-Eco1 clustered in clade 1, whereas toxic RcaT-Sen2 clustered in clade 3. Interestingly, although tetramerisation appears architecturally conserved within clades 2 and 3, conservation of the predicted interfaces at the sequence level is low (Extended Data Fig. 8e). We next sought to experimentally validate a subset of the oligomeric state predictions.

We first examined all remaining RcaTs with previously characterised toxicity profiles^6^ (Fig. 4c, bold letters). RcaT-Eco3 (Fig. 4b, clade 1) is non-toxic^6^ and could be purified in its wild-type form, eluting exclusively as a dimer by SEC (Fig. 4a). RcaT-Eco9 and RcaT-Eco10 (Fig. 4b, both clade 3) are toxic^6^ and required catalytic inactivation for expression. Catalytically inactive RcaT-Eco10 formed oligomers even exceeding tetramers (Fig. 4a). In contrast, catalytically inactive RcaT-Eco9 could not be purified, so we used mutagenesis to test its tetramer model. The predicted RcaT-Eco9 dimer–dimer interface is similar to that of RcaT-Sen2, with Eco9 S254 corresponding to Sen2 S253 (Fig. 4d; Fig. 3c). Introducing the S254D substitution in RcaT-Eco9 abolished toxicity (Fig. 4c, boxed), mirroring the effect of the S253G mutation in Sen2 (Fig. 3d). This supports a role for the predicted dimer–dimer interface in RcaT-Eco9 toxicity. Together, these results validate the AlphaFold-based oligomeric predictions and establish a strong correlation between RcaT dimer–dimer assembly and intrinsic toxicity.

We then tested whether toxicity could be predicted *a priori* from oligomerisation propensity. Among the remaining RcaTs, only RcaT-Vch3 (Fig. 4b, clade 1) had been partially characterised^25^, but its toxicity was unknown. Spot assays revealed that RcaT-Vch3 is non-toxic, in agreement with its predicted dimeric state (Fig. 4c). We then selected four additional RcaTs spanning the phylogeny and assigned nomenclature^26^: RcaT-Hin1 (Fig. 4b, clade 2), RcaT-Vvu1 (Fig. 4b, clade 3), RcaT-Cle1 and RcaT-Bth2 (Fig. 4b, both clade 1). AlphaFold confidently predicted tetramers for RcaT-Hin1 and RcaT-Vvu1, but not for RcaT-Cle1 or RcaT-Bth2 (Fig. 4b). Spot assays confirmed that the tetramer-predicted RcaTs were toxic, whereas the dimer-predicted RcaTs were non-toxic (Fig. 4c).

Together, phylogeny-resolved AlphaFold predictions, biochemical analysis, and functional validation establish oligomerisation beyond the dimer as a defining and predictive hallmark of RcaT toxicity. Although all RcaT retrons defend against phage infection^6^, only retrons carrying RcaTs that can oligomerise beyond a dimer are directly toxic to the host.

## Discussion

This study provides a structural view of Retron-Sen2 and reveals how single-site msDNA methylation triggers complex disassembly to activate phage defence. In the Retron-Sen2 toxin–antitoxin complex, the antitoxin sequesters RcaT as a dimer, occluding its dimer–dimer interface. Further oligomerisation of RcaT through this interface is essential for its toxicity, hence explaining why it remains inactive in the toxin–antitoxin complex. We show that methylation disrupts the RcaT–msDNA interaction and promotes complex disassembly. This likely occurs by a second RcaT dimer binding to the previously occluded interface and outcompeting RT for overlapping binding sites, enabling dimer–dimer assembly (Fig. 4e). A possible route to complete release of RcaT is outlined in Extended Data Fig. 9.

DNA methyltransferases have been implicated in the activation of diverse type II retrons^6,18,19^. Retron-Eco9 may share the activation mechanism we report for Retron-Sen2, as their RcaTs are similar in toxicity^6^ and oligomeric state. By contrast, how methylation activates Retron-Eco1 is more puzzling. Its reported methylation site lies far from RcaT and is mismatched on the complementary strand^18^, making disassembly driven by steric hindrance unlikely. Even if methylation displaced msDNA, RcaT-Eco1 cannot oligomerise further to outcompete the RT. Moreover, RcaT-Eco1 is non-toxic^6^, so its release would be functionally inconsequential. Other retron complexes have also been shown to disassemble upon triggering^7,10,17^. In type I Retron-Eco4^10^ and -Eco7^17^ and type IX Retron-Yin1^7^, nuclease-mediated msDNA degradation triggers effector release^10,17^, but a requirement for an additional activation step analogous to RcaT dimer–dimer assembly has not been established. Together, these observations suggest that retron activation is mechanistically diverse at multiple levels. The same trigger can promote distinct pathways even within the same retron type, while different retron types can converge on complex disassembly as an activation mechanism yet differ in how effector release is coupled to downstream activity.

Beyond its role in toxin–antitoxin complex disassembly, dimer–dimer assembly is also essential for toxicity. This extends beyond RcaT-Sen2 and emerges as a defining property of toxic RcaTs, whereas non-toxic homologues appear intrinsically dimeric. This requirement for dimer–dimer assembly provides a mechanistic explanation for previously unexplained observations. The S253G mutation in RcaT-Sen2, previously identified as a suppressor of toxicity^6^, is shown here to disrupt the dimer–dimer interface. Similarly, RcaT-Eco9 was recently shown to be disarmed by T7 kinase as a phage counter-defence mechanism^27^. One of the phosphorylated residues is S254^28^, which corresponds to S253 in RcaT-Sen2. Our finding that a phosphomimetic S254D mutation abolishes RcaT-Eco9 toxicity provides a structural rationale, linking this counter-defence modification to inhibition of dimer–dimer assembly. Thus, the same oligomeric switch that governs retron activation and RcaT toxicity also appears to constitute a vulnerability exploited by phage counter-defence.

Despite the mechanistic insights provided by our study, several questions remain. Retron-Sen2 can form filaments through interactions mediated by the retained msr fragment, an assembly mechanism not previously described in other retrons. Filament formation itself is also observed in the homologous Retron-Eco1^13,18^, although its architecture is reversed relative to Retron-Sen2. Similar assemblies are increasingly observed across bacterial defence systems^7,29–32^, yet whether they form *in vivo* and what physiological roles they serve remain unclear. The role of filament formation is not resolved by our current model, as all features of Retron-Sen2 inhibition and activation can be explained by the core toxin–antitoxin complex. Another unresolved question is why dimer–dimer assembly is required for RcaT toxicity despite being seemingly dispensable for its catalytic activity. One possibility is that it enables toxic RcaTs to ADP-ribosylate an essential host target, although the identity of this target and the underlying mechanism remain to be established. Oligomerisation also does not explain condition-dependent activity among toxic RcaTs^6^. RcaT-Sen2 is toxic only below 30 °C or under anaerobic conditions, whereas RcaT-Eco9 is constitutively toxic^6^. These open questions motivate future studies to define how RcaT acts on its physiological targets, uncover the mechanisms regulating its conditional activity and determine how retron filament formation contributes to phage defence.

## Methods

### Plasmid construction

Plasmids encoding native retrons Sen2, Eco1, Eco3, Eco9, Eco10, and Vch3 were obtained from Bobonis *et al.*^6^ and used to subclone individual components of these systems. RcaT homologues from retrons Hin1, Vvu1, Cle1, and Bth2 were synthesised and codon-optimised for *E. coli* expression by GenScript (Piscataway, NJ, USA) and similarly subcloned. Complete details of plasmid construction and site-directed mutagenesis are provided in Supplementary Data 5. Plasmids were transformed into chemically competent *E. coli* DH5α, and transformants were selected using appropriate antibiotics. All plasmids were purified using the NucleoSpin Plasmid kit (Macherey-Nagel, Düren, Germany) and verified by sequencing at Microsynth AG (Balgach, Switzerland) or Eurofins Genomics (Ebersberg, Germany).

### Protein expression and purification

For expression of Retron-Sen2, plasmids pPS45 and pPS56 were co-transformed into *E. coli* BL21(DE3) (Novagen, Merck KGaA, Darmstadt, Germany). A 6 L culture was grown in LB medium supplemented with kanamycin (50 μg mL⁻¹) and spectinomycin (50 μg mL⁻¹) at 37 °C with shaking until reaching an OD₆₀₀ of 1.0. The culture was then cooled to 16 °C and induced with 0.3 mM IPTG for 18 h. Cells were harvested by centrifugation (20 min, 6,600 × g).

The purification protocol was adapted and optimised from Wang *et al.*^12^. The cell pellet was resuspended in 200 mL lysis buffer (40 mM Bis-Tris, 300 mM NaCl, pH 6.6). Cells were lysed using an emulsifier at 20,000 psi (three passes). The lysate was clarified by centrifugation (20 min, 41,200 × g) and filtered through a 0.22 µm membrane. The cleared lysate was incubated for 1 h with 5 mL Strep-Tactin XT 4Flow resin (IBA Lifesciences, Göttingen, Germany), pre-equilibrated in lysis buffer, then loaded onto a gravity flow column. After washing with 300 mL lysis buffer, bound proteins were eluted in 20 mL elution buffer (40 mM Bis-Tris, 450 mM NaCl, 50 mM biotin, pH 6.6). The eluate was immediately applied to a Mono Q 5/50 GL anion exchange column (Cytiva, Marlborough, MA, USA), connected to an ÄKTA Pure system (Cytiva, Marlborough, MA, USA) and pre-equilibrated with lysis buffer. Protein was eluted using a 100 mL linear gradient (300–800 mM NaCl in 40 mM Bis-Tris, pH 6.6). Peak fractions corresponding to Retron-Sen2 were pooled, concentrated to 500 µL, and subjected to SEC on a Superdex 200 Increase 10/300 GL column (Cytiva, Marlborough, MA, USA), pre-equilibrated with SEC buffer (20 mM Bis-Tris, 250 mM NaCl, pH 6.6), again using the ÄKTA Pure system. Protein integrity was confirmed by SDS–PAGE. The final sample was concentrated to 0.8 mg mL⁻¹, as determined by BCA assay. The Retron-Sen2 antitoxin was purified using the same protocol.

For expression of RcaT-Sen2 E107Q, plasmid pPS22 was transformed into *E. coli* BL21(DE3) LOBSTR^33^ (Kerafast, Boston, MA, USA). A 2 L culture was grown in LB medium supplemented with spectinomycin (50 μg mL⁻¹) at 37 °C with shaking until reaching an OD₆₀₀ of 0.6. The culture was then cooled to 15 °C and induced with 0.3 mM IPTG for 48 h. Cells were harvested by centrifugation (20 min, 6,600 × g).

The cell pellet was resuspended in 80 mL lysis buffer (40 mM Bis-Tris, 1 M NaCl, 40 mM imidazole, pH 6.6). Cells were lysed using an emulsifier at 20,000 psi (three passes). The lysate was clarified by centrifugation (20 min, 41,200 × g) and filtered through a 0.22 µm membrane. The cleared lysate was incubated for 1 h with 2 mL Protino Ni-NTA resin (Macherey-Nagel, Düren, Germany), pre-equilibrated in lysis buffer, and then loaded onto a gravity flow column. The column was washed with 120 mL lysis buffer followed by 30 mL wash buffer (40 mM Bis-Tris, 1 M NaCl, 100 mM imidazole, pH 6.6), and eluted in 10 mL elution buffer (40 mM Bis-Tris, 1 M NaCl, 500 mM imidazole, pH 6.6). EDTA was then added to the eluate to a final concentration of 10 mM, along with 1 mL His-tagged TEV protease (1 mg mL⁻¹), and the mixture was dialysed overnight against dialysis buffer (40 mM Bis-Tris, 1 M NaCl, pH 6.6). The dialysed eluate was incubated with 1 mL Protino Ni-NTA resin pre-equilibrated in dialysis buffer, then loaded onto a gravity flow column. The flowthrough was collected, concentrated to 500 μL, and subjected to SEC on a Superdex 200 Increase 10/300 GL column, pre-equilibrated with SEC buffer (20 mM Bis-Tris, 500 mM NaCl, pH 6.6), using an ÄKTA Pure system. Protein integrity was confirmed by SDS–PAGE and BN-PAGE, and the final sample was concentrated to 1.5 mg mL⁻¹, as determined by Nanodrop. All other RcaT variants and homologues were purified using a similar protocol.

### Isolation and digestion of msDNA

One volume of freshly purified Retron-Sen2 complex was mixed with one volume of phenol:chloroform:isoamyl alcohol (25:24:1), vortexed for 3 seconds, and centrifuged at 14,000 × g for 2 minutes at room temperature. The aqueous phase was collected and combined with 2 volumes of 100% ethanol and 0.1 volume of 3 M sodium acetate, followed by incubation at –70 °C for 1 hour. The sample was then centrifuged at 14,000 × g for 5 minutes at 4 °C. After discarding the supernatant, the pellet was rinsed with 70% ethanol and centrifuged again at 14,000 × g for 3 minutes at 4 °C. The supernatant was removed, and the pellet was air-dried before resuspension in nuclease-free water. To assess its composition, aliquots of the isolated msDNA were treated separately with DNase I and RNase A, and analysed alongside an untreated control on a 15% TBE–urea polyacrylamide gel.

### *In vitro* methylation assay

Retron-Sen2 complex was purified as previously described, with the exception that the SEC buffer was replaced with methylation buffer (40 mM Tris-HCl, 50 mM NaCl, pH 7.4). The purified complex was divided into two equal aliquots, and S-adenosylmethionine (SAM; New England Biolabs, Ipswich, MA, USA) was added to a final concentration of 160 μM. DNA adenine methyltransferase (Dam; New England Biolabs, Ipswich, MA, USA) was added to one aliquot, while the other received heat-inactivated Dam (65 °C for 20 minutes) as a control. The samples were incubated for 1 hour at 37 °C. msDNA was isolated from a small portion of each methylated and unmethylated reaction as previously described and digested with either MboI or DpnI (New England Biolabs, Ipswich, MA, USA) to confirm complete methylation or its absence. A second portion of each sample was used for RcaT pull-down via the RT affinity tag. Finally, a third portion was analysed by SEC in SEC buffer (20 mM Bis-Tris, 250 mM NaCl, pH 6.6) to assess complex disassembly.

### ADP-ribosylation assay

RcaT-Sen2 E107Q (catalytically inactive negative control) and R301D were purified as previously described and incubated with 1 mM NAD⁺ for 30 minutes. Proteins at 0.1 mg mL^−1^ were heated at 70 °C for 3 minutes in SDS loading buffer and resolved by SDS–PAGE, followed by transfer to a PVDF membrane. Membranes were blocked with 5% BSA in TBST for 1 hour at room temperature and incubated overnight at 4 °C with anti-Poly/Mono-ADP Ribose (D9P7Z) Rabbit mAb (Cell Signaling Technology, Danvers, MA, USA) primary antibody. After three washes, membranes were incubated for 1 hour at room temperature with goat anti-rabbit IgG (F(abʹ)₂ fragment), HRP-conjugated secondary antibody (Merck, Darmstadt, Germany). Blots were developed using SuperSignal™ West Pico PLUS Chemiluminescent Substrate (Thermo Fisher Scientific, Waltham, MA, USA) and imaged.

### Mass photometry

Mass photometry measurements were performed on a TwoMP mass photometer (Refeyn Ltd., Oxford, UK). A clean coverslip was placed on the sample holder, and 18 μL of SEC buffer (20 mM Bis-Tris, 250 mM NaCl, pH 6.6) was added for focusing, followed by 1 μL of Retron-Sen2 antitoxin at 250 nM. Mass contrast calibration was conducted using the NativeMark unstained protein standard (Thermo Fisher Scientific, Waltham, MA, USA) prior to data export. Calibrated movies were analysed using Photomol^34^, with Gaussian fitting and binning applied before final visualisation in R version 4.3.3 (R Core Team, 2024).

### NanoDSF

RcaT–Sen2 E107Q was diluted in SEC buffer (20 mM Bis-Tris, 500 mM NaCl, pH 6.6) to a final concentration of 0.3 mg mL^⁻¹^. Two capillaries containing 10 µL of sample each were loaded into a Prometheus nanoscale differential scanning fluorimeter (NanoTemper, Munich, Germany). A temperature ramp from 20 °C to 95 °C was applied. Tm fitting was performed using MoltenProt^35^ and visualised in R version 4.3.3.

### Cryo-EM sample preparation and data collection

For all three complexes, 5 μL of freshly purified sample was applied to grids that had been glow-discharged at 25 mA for 120 s using a GloQube device (Quorum Technologies Ltd., Laughton, East Sussex, UK). For Retron-Sen2, the sample was at a concentration of 0.8 mg mL⁻¹ and applied to Quantifoil R2/1, Cu 300 mesh grids (Jena Bioscience, Jena, Germany). The Retron-Sen2 antitoxin was applied at the same concentration to Quantifoil R1.2/1.3, Cu 300 mesh grids (Jena Bioscience). RcaT-Sen2 was prepared at a concentration of 1.5 mg mL⁻¹ and applied to Quantifoil R2/1, Cu 300 mesh grids. After sample application, grids were incubated for 10 s and blotted from both sides using a Vitrobot Mark IV (Thermo Fisher Scientific, Waltham, MA, USA) at 100% humidity and 4 °C, with blot force −7 and a blot time of 2.5 s. Finally, they were plunge-frozen in a 63:37 propane-ethane mixture.

Cryo-EM data were acquired on a Titan Krios transmission electron microscope (Thermo Fisher Scientific, Waltham, MA, USA) operated at 300 kV, equipped with a K3 direct electron detector and a GIF Quantum energy filter with a 20-eV slit width (Gatan, Pleasanton, CA, USA). Movies were recorded in counting mode using EPU software (Thermo Fisher Scientific, Waltham, MA, USA) at a nominal magnification of ×130,000, corresponding to a calibrated pixel size of 0.68 Å, and a defocus range of −0.75 to −2.5 μm. For the Retron-Sen2 complex, 10,228 movies were collected, each fractionated over 50 frames with a total dose of 58 e⁻/Å². For the Retron-Sen2 antitoxin, 3,637 movies were collected, also fractionated over 50 frames, with a total dose of 51.1 e⁻/Å². For RcaT-Sen2, 5,517 movies were collected, each fractionated over 40 frames with a total dose of 46.36 e⁻/Å².

### Cryo-EM data processing

All datasets were processed using cryoSPARC v4.4.1^36^. Patch Motion Correction was used to estimate and correct for full-frame drift and local sample deformation, followed by Patch Contrast Transfer Function (CTF) Estimation to fit local CTF models to each micrograph. Micrographs with estimated CTF resolutions worse than 5 Å were manually excluded. Initial particle picking was performed either on full or partial datasets using the Blob Picker. Extracted particles underwent multiple rounds of 2D classification, and resulting high-quality class averages were used as input for the Template Picker to enable full-dataset particle picking.

For Retron-Sen2, initial 2D classification of Blob Picker–derived particles revealed structural heterogeneity, distinguishing between the toxin–antitoxin complex and filament assemblies. The corresponding 2D classes were separated and used to generate two dedicated Template Pickers. For the Retron-Sen2 toxin–antitoxin complex, particles were extracted using a 448-pixel box size. After three rounds of 2D classification, *ab initio* reconstruction was performed into two classes. The better-resolved class was selected for homogeneous refinement in C1, followed by non-uniform (NU) refinement^37^ in C1 and finally C2, yielding a high-resolution map at 3.37 Å. For the Retron-Sen2 filament, particles were extracted with a 720-pixel box size and Fourier-cropped to 540 pixels to facilitate processing. Five rounds of 2D classification were performed, retaining only distinct views of 2-unit filaments. These particles were subjected to *ab initio* reconstruction into two classes, and the better-resolved class was selected for homogeneous refinement in C1. A subsequent 3D classification into six classes allowed exclusion of junk particles. Non-uniform refinement was performed in two stages: first in C1, then in C2, yielding a reconstruction of the 2-unit filament at 3.55 Å. To focus on the central filament interface, local refinement using a soft mask covering this region produced a 3.29 Å map.

For the Retron-Sen2 antitoxin, particles were extracted using a 256-pixel box size. Four rounds of 2D classification revealed structural heterogeneity, stemming from flexibility in the msd. Two states were particularly prominent: one resembling the antitoxin conformation observed when bound to RcaT, and another in which the msd adopted a different orientation. The corresponding 2D classes were separated. For the first state we performed *ab initio* reconstruction into two classes, selected the better-resolved class, and proceeded with homogeneous reconstruction followed by local refinement in C1 using a soft mask that excluded the tip of the msd, yielding a 4.99 Å map. For the second state, we performed *ab initio* reconstruction with two classes and selected the best-resolved particles. To capture more of the differently oriented msd, particles were re-extracted using a 360-pixel box size and subjected to a second round of *ab initio* reconstruction, followed by NU refinement in C1. The resulting map (5.57 Å) suffered from a limited number of particles and orientation bias.

For the antitoxin–free RcaT-Sen2, initial 2D classification of Blob Picker–derived particles again revealed structural heterogeneity. The corresponding 2D classes were separated and used to generate two dedicated Template Pickers. For the tetramer, particles were extracted with a 480-pixel box size. After three rounds of 2D classification, *ab initio* reconstruction into two classes was performed. The resulting volumes were used twice as templates for a four-class heterogeneous refinement in C1. The best-resolved class was selected and subjected to NU refinement in C1, followed by NU refinement in C2. The particles were then symmetry-expanded in C2 and locally refined in C1 using a global mask, yielding a 3.19 Å map. For higher-order oligomers, particles were extracted with a 600-pixel box size and Fourier-cropped to 540 pixels to facilitate processing. Multiple rounds of 2D classification were performed, and the selected particles were subjected to two-class *ab initio* reconstruction. The better-resolved class volume was used as a template for NU refinement in C1, while retaining particles from both classes. Subsequent NU refinement in C2 produced a 3.78 Å map of a decamer. Local refinement with a soft mask covering a hexamer within the decamer improved the resolution to 3.71 Å.

In all cases, resolution was estimated from Gold-standard Fourier shell correlation (GSFSC) curves using the 0.143 criterion. Local resolution was assessed, and maps were sharpened with DeepEMhancer^38^ for visualisation.

### Model building

The models for the Retron-Sen2 toxin–antitoxin complex and filament were built iteratively, using the best-resolved regions from each map to model the corresponding subunits. For the toxin–antitoxin complex, an AlphaFold3^39^ prediction of the protein components was manually fit into the C2 map using ChimeraX 1.7.1^40^ and refined in ISOLDE^41^ using self-restraints. The refined model was then used as input for gapTrick^23,24,42,43^, an AlphaFold2-based approach, to generate an improved structure, which was further refined in ISOLDE using self-restraints. The msd stem was modelled as canonical B-form DNA in Coot^44^, with the msd loop and ssDa regions also built *de novo*. The full assembly was then refined in Coot using self-restraints at a cut-off of 5 Å. This model notably lacked ssDb and the entire msr. Separately from the toxin–antitoxin complex, an AlphaFold3 model comprising the RT, DNA–RNA duplex, and RSLb (Extended Data Fig. 2d) was fit into the C2 local refinement map of the filament interface using ChimeraX. The ssDb region preceding the DNA–RNA duplex and the ssRb segment following RSLb were built *de novo* in Coot and refined alongside the predicted antitoxin model using self-restraints at a cut-off of 5 Å. Because the filament interface model lacked RcaT, the DSL and ssDa, these components were copied from the toxin–antitoxin complex model and manually fit into the corresponding density using ChimeraX. Any regions lacking supporting density were deleted. The resulting filament interface model was refined in ISOLDE with all double-stranded nucleic acids pinned, while self-restraints were applied to the protein components. The refined ssDb, RSLb, ssRb, and DNA– RNA duplex from the filament interface model were incorporated into the toxin–antitoxin complex model and fit to its map. This composite model was then refined in ISOLDE, again pinning double-stranded nucleic acids and applying self-restraints to the protein. We chose not to deposit a PDB model for the 2-unit filament, only its corresponding map in the EMDB, due to a significant decline in map quality with increasing distance from the filament interface. To approximate the two-unit filament shown in Extended Data Fig. 4a, we performed rigid-body fitting of two toxin–antitoxin complexes into the two-unit cryo-EM map. The four-unit Retron-Sen2 filament model shown in Extended Data Fig. 4d was generated by successive superposition of overlapping units of the two-unit model. Both models are provided in Supplementary Data 6.

For the Retron-Sen2 antitoxin, we took one antitoxin copy from the filament junction model and fit it into the 4.99 Å cryo-EM map. Minimal adjustments were made during refinement in Coot using self-restraints set to a cut-off of 5 Å. We did not build a refined atomic model of the msd 90° rotation state because of the limited map quality, but an approximate model used only for illustrative purposes in Fig. 4e is provided in Supplementary Data 6.

For antitoxin-free RcaT-Sen2, AlphaFold-Multimer predictions of the tetramer and hexamer were manually fitted into the corresponding C2 maps using ChimeraX and refined in ISOLDE with self-restraints. The refined models were then processed with gapTrick, and the improved structures were further refined in ISOLDE with self-restraints. In the case of the hexamer, flexible regions lacking supporting density were deleted.

All models were refined against their respective maps using Phenix real-space refinement^45,46^ with secondary structure restraints and validated using Phenix’s comprehensive validation tool^46,47^. Validation statistics are shown in Extended Data Table 1. All model visualisation and analysis were carried out in ChimeraX, PyMOL and PDBePISA^48^.

### Spot assays

For spot growth assays, plasmids were transformed into *E. coli* BW25113. Single bacterial colonies were inoculated in 5 mL LB plus antibiotic and incubated overnight at 37 °C in a roller drum. Saturated overnight cultures (OD595 ∼6) were serially diluted (10-fold steps) seven times (undiluted to 10-7) in LB. ∼10 μL of culture dilutions were spotted using a 96-pinner (V&P Scientific, VP 404) on Singer LB plates containing the appropriate antibiotic and/or arabinose, when applicable. Plates were incubated at 37 °C for 12-14 hours before images were taken.

### Phylogenetic tree construction and oligomeric state predictions

Sequences of 5,938 RcaTs used to construct the phylogenetic tree were obtained from Bobonis *et al.*, and the tree was reconstructed following the methodology described in the original publication^6^. Briefly, sequences were clustered at 99% identity using CD-HIT^49^, resulting in 776 unique entries. These sequences were aligned with MAFFT^50^ using default parameters, and maximum-likelihood phylogenies were inferred with FastTree2^51^. Sequence-level conservation was visualised by ConSurf^52,53^.

For the smaller, representative tree, 32 RcaT sequences were selected from across the full tree (Extended Data Fig. 8c). These were aligned using MAFFT with the L-INS-i strategy (--lop 3.7), and phylogenies were inferred as above. Each sequence was subsequently used to predict both homodimeric and homotetrameric assemblies using AlphaFold2 version 2.3.2 with a jackhmmer-based^54^ MSA generation pipeline and default parameters, generating 25 models per run. The resulting ipTM scores were plotted alongside the tree. Visualisation was performed using R version 4.3.3.

AlphaFold pTM and ipTM scores for the homodimeric and homotetrameric predictions of the selected RcaTs are available in Supplementary Data 2. Full dimer and tetramer predictions (PAE, sequence coverage, and contact plots, as well as PDB coordinates) are provided in Supplementary Data 3 and 4, respectively.

## Supplementary Information

Supplementary Data 1 | PISA analysis of interfaces in structures determined in this study

Supplementary Data 2 | Selected RcaT AlphaFold pTM and ipTM annotations

Supplementary Data 3 | AlphaFold prediction data for the RcaT homodimers

Supplementary Data 4 | AlphaFold prediction data for the RcaT homotetramers

Supplementary Data 5 | Plasmids and primers used in this study

Supplementary Data 6 | Approximate structural models used only for illustrative purposes

## Data Availability

The coordinates of the cryo-EM structures of the Retron-Sen2 toxin–antitoxin complex, filament interface, antitoxin, RcaT tetramer, and hexamer have been deposited in the Protein Data Bank (PDB) under accession codes 34PA, 34PL, 34PF, 34PB, and 34PC, respectively. The corresponding density maps have been deposited in the Electron Microscopy Data Bank (EMDB) under accession codes EMD-100081, EMD-100090, EMD-100088, EMD-100082, and EMD-100083. Additional maps for the Retron-Sen2 two-unit filament, the alternative antitoxin conformation, and decameric RcaT have been deposited in the EMDB under accession codes EMD-100080, EMD-100079, and EMD-100084 respectively.

## Source Data

Source Data 1 | Uncropped images

Source Data 2 | Plot values

## Acknowledgments

We thank the EM facility at the Centre for Structural Systems Biology, Hamburg, for access to instrumentation and support with sample screening and data collection. We also thank the SPC facility at EMBL Hamburg and the Proteomics Core Facility at EMBL Heidelberg for access and technical support. We acknowledge the use of the HPC infrastructure at EMBL Heidelberg. We are grateful to Sihyun Sung (EMBL Grenoble), Yehudi Bloch (DESY, Hamburg), Sebastian Eustermann (EMBL Heidelberg), Katherine Beckham (University of Newcastle), Sarah J. Coulthurst (University of Dundee), and Genady Pankov (University of Dundee) for their technical and scientific advice at various stages of the project. This work was supported by core funding from the European Molecular Biology Laboratory.

## Author contributions

A.T., J.B., and P.S. conceived the study. A.T. and M.W. supervised and supported the project. A.L.J.Y., D.K., F.C., and P.S. constructed plasmids. P.S. performed and analysed biochemical and biophysical experiments. E.M. and P.S. prepared grids, performed screening, collected and processed cryo-EM data. P.S. carried out integrative modelling with input from G.C. M.W. and P.S. interpreted the structures. A.L.J.Y. and D.K. performed the microbiology experiments. P.S. performed the phylogenetic analysis, and G.C. and P.S. carried out AlphaFold predictions for annotation of the tree. P.S. generated figures. M.W., A.T., and P.S. wrote the manuscript with input from all authors. All authors have read and approved the manuscript.

## Extended Data Figures

**Extended Data Figure 1.**
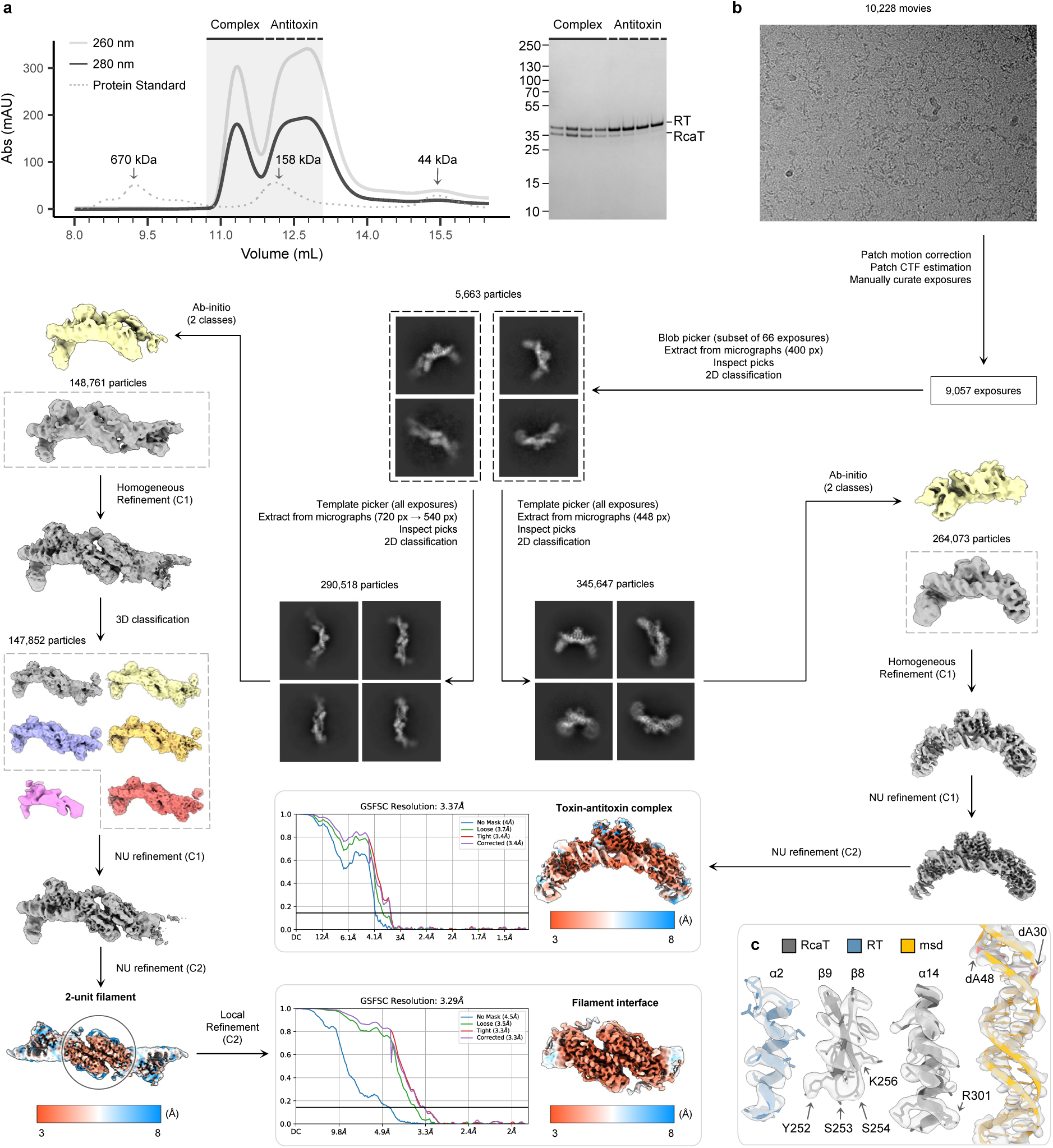
Structural characterisation of Retron-Sen2. **a,** SEC resolves Retron-Sen2 into two peaks. The toxin– antitoxin complex elutes earlier, whereas the later peak contains predominantly free antitoxin. Light grey, dark grey and dotted traces indicate UV absorbance at 260 nm, 280 nm and protein standards, respectively. An SDS–PAGE gel of fractions from both peaks is shown at right. **b**, Cryo-EM data processing. The Retron-Sen2 toxin–antitoxin complex was reconstructed to an estimated resolution of 3.37 Å. A two-unit filament was also reconstructed, with local refinement of the filament interface yielding a map with an estimated resolution of 3.29 Å. GSFSC curves and local resolution maps are shown for both reconstructions. **c**, Close-ups show selected regions of the cryo-EM map (transparent surface) with the fitted atomic model. These regions mediate RcaT inhibition by the antitoxin. RT helix α2 (blue) interacts with RcaT; the RcaT β8–β9 hairpin contacts the msDNA methylation site, while RcaT helix α14 contacts both the methylation site and RT. The msd (orange) is central to RcaT inhibition, with the methylation site shown in red. Key residues near the methylation site are highlighted.

**Extended Data Figure 2.**
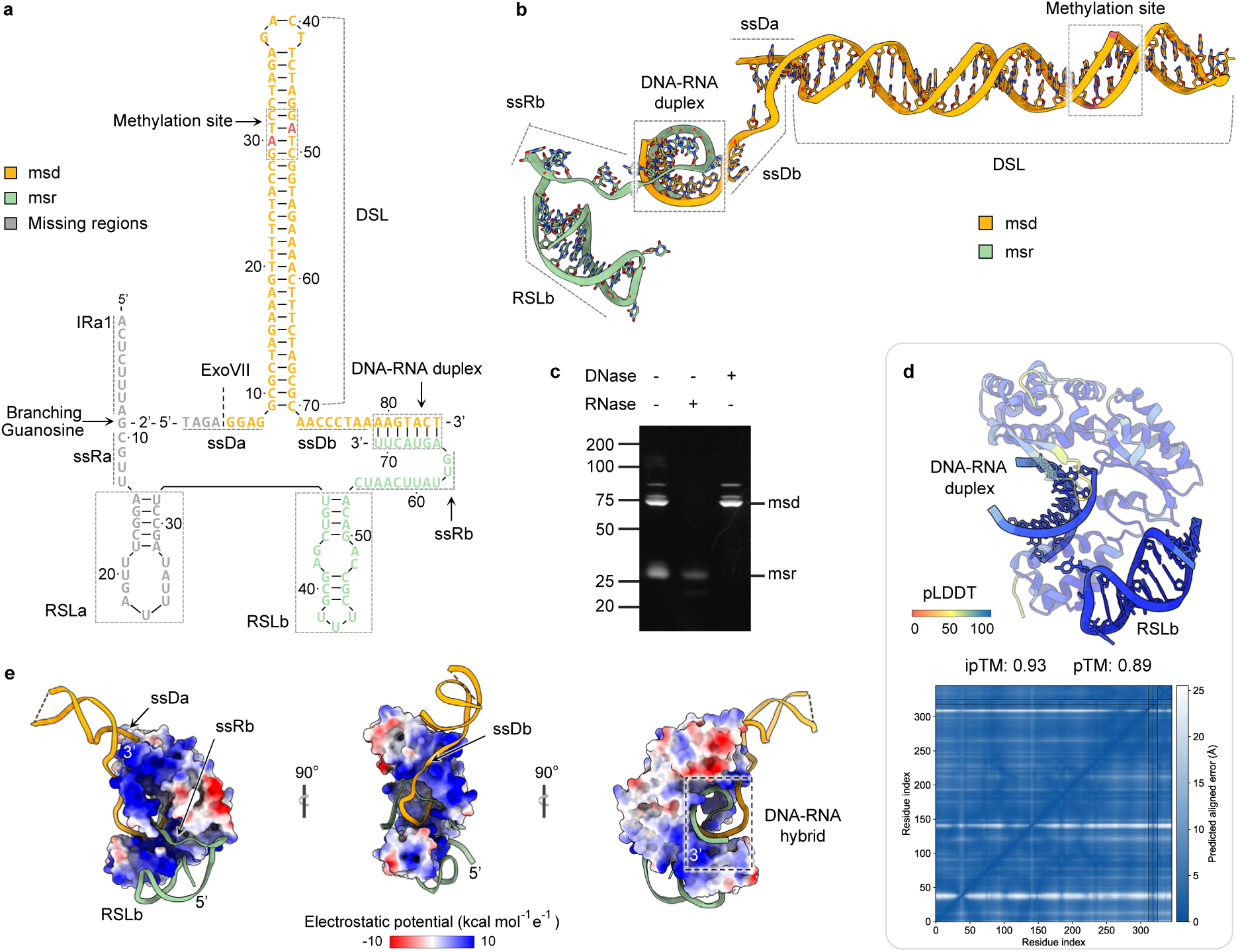
Retron-Sen2 retains a fragment of msr after ExoVII processing. **a,** A schematic shows msd (orange) and msr (green) in processed Retron-Sen2 msDNA. Nucleotides of msr not present in the cryo-EM structures are shown in grey. ExoVII processing appears to have occurred, releasing the first four nucleotides of msd and the first 32 nucleotides of msr. Unexpectedly, the final 39 nucleotides of msr persist, likely protected through interactions with RT, whereas the remaining regions are likely susceptible to host RNases. Retention of an msr fragment has recently been observed in other ExoVII-processed retrons^8,55^. Structural elements are annotated as follows: DSL, DNA stem–loop; ssDa and ssDb, single-stranded DNA a and b; RSLa and RSLb, RNA stem–loops a and b; ssRa and ssRb, single-stranded RNA a and b; IRa, inverted repeat a. **b**, Atomic model of the processed msDNA, comprising msd and msr. **c**, msDNA resolves into two predominant nucleic acid species on a denaturing TBE–urea gel: the upper band is DNase-sensitive, whereas the lower band is RNase-sensitive. **d**, AlphaFold3 predicts a complex comprising RT, RSLb and the DNA–RNA duplex with high confidence. The model is coloured by pLDDT confidence scores, and the predicted aligned error (PAE) plot is shown below. This prediction guided building of the antitoxin model (see Methods). **e**, The RT engages msr, the DNA–RNA duplex, ssDa and ssDb through an extended positively charged surface. The RT surface is coloured by electrostatic potential according to the scale shown below.

**Extended Data Figure 3.**
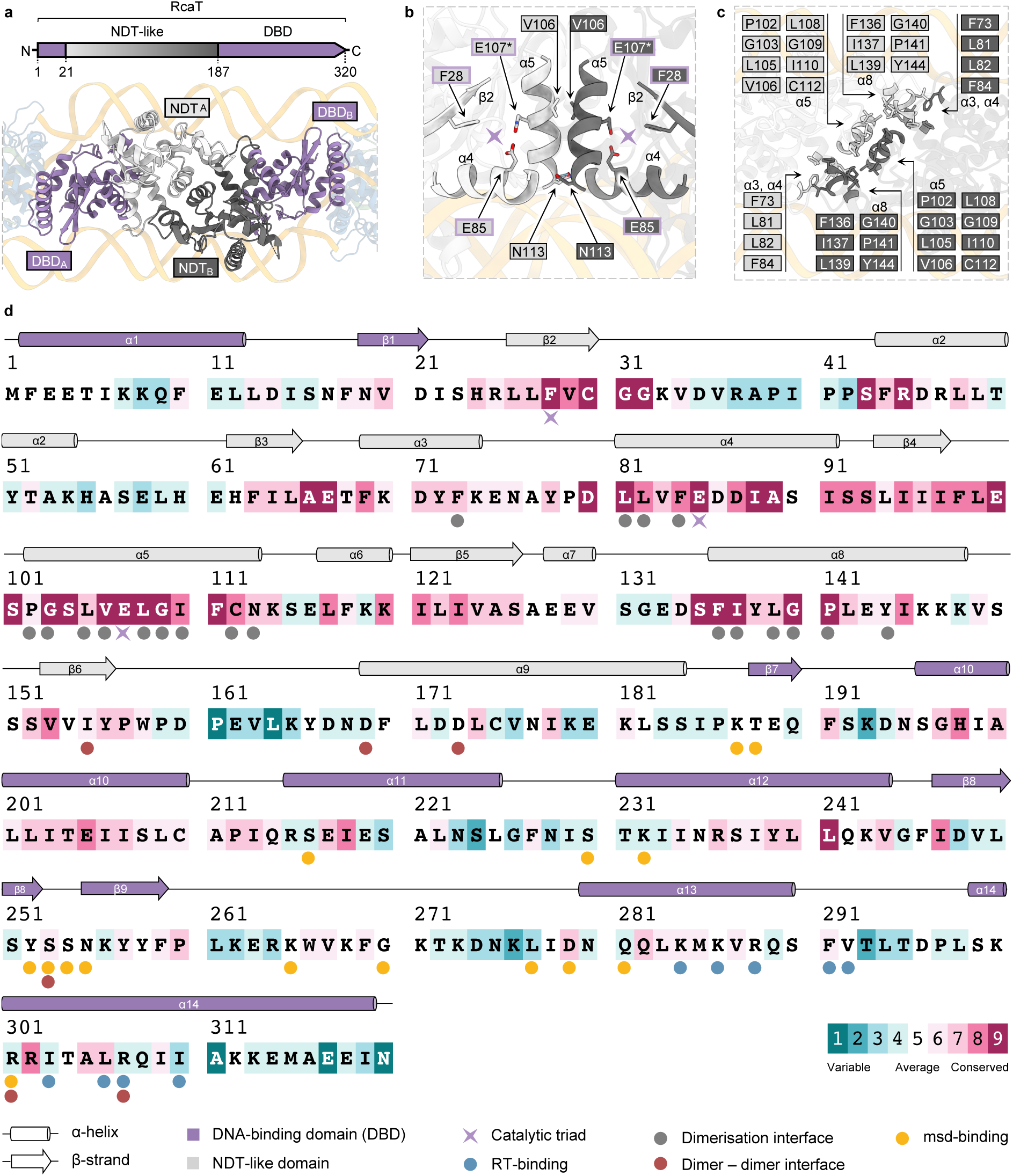
RcaT dimerises through a conserved hydrophobic interface within its NDT-like domain. **a,** RcaT-Sen2 forms a symmetric dimer via its NDT-like domain (grey; residues 22-186), with DNA-binding domains (DBDs; purple; residues 1-21 and 187-320) projecting outward. **b**, The NDT active sites are located at the centre of the dimer (lilac asterisks). The catalytic triad comprises F28, E85 and E107 (outlined in lilac; E107* was mutated to Q for structure determination). V106 and N113 contribute to the dimerisation interface. **c**, Dimerisation is primarily mediated by a hydrophobic core of 18 residues per chain; contributing residues are indicated. **d**, ConSurf analysis shows RcaT-Sen2 conservation across the RcaT phylogeny, based on a multiple sequence alignment of 776 unique RcaTs represented in the phylogenetic tree of Extended Data Fig. 8c. Default ConSurf colour coding indicates sequence-level conservation. Domain-coloured secondary-structure elements are annotated above the sequence. Residues involved in catalytic activity, dimerisation, dimer–dimer assembly, RT binding and msd binding are indicated below the sequence. All annotations are explained in the bottom inset.

**Extended Data Figure 4.**
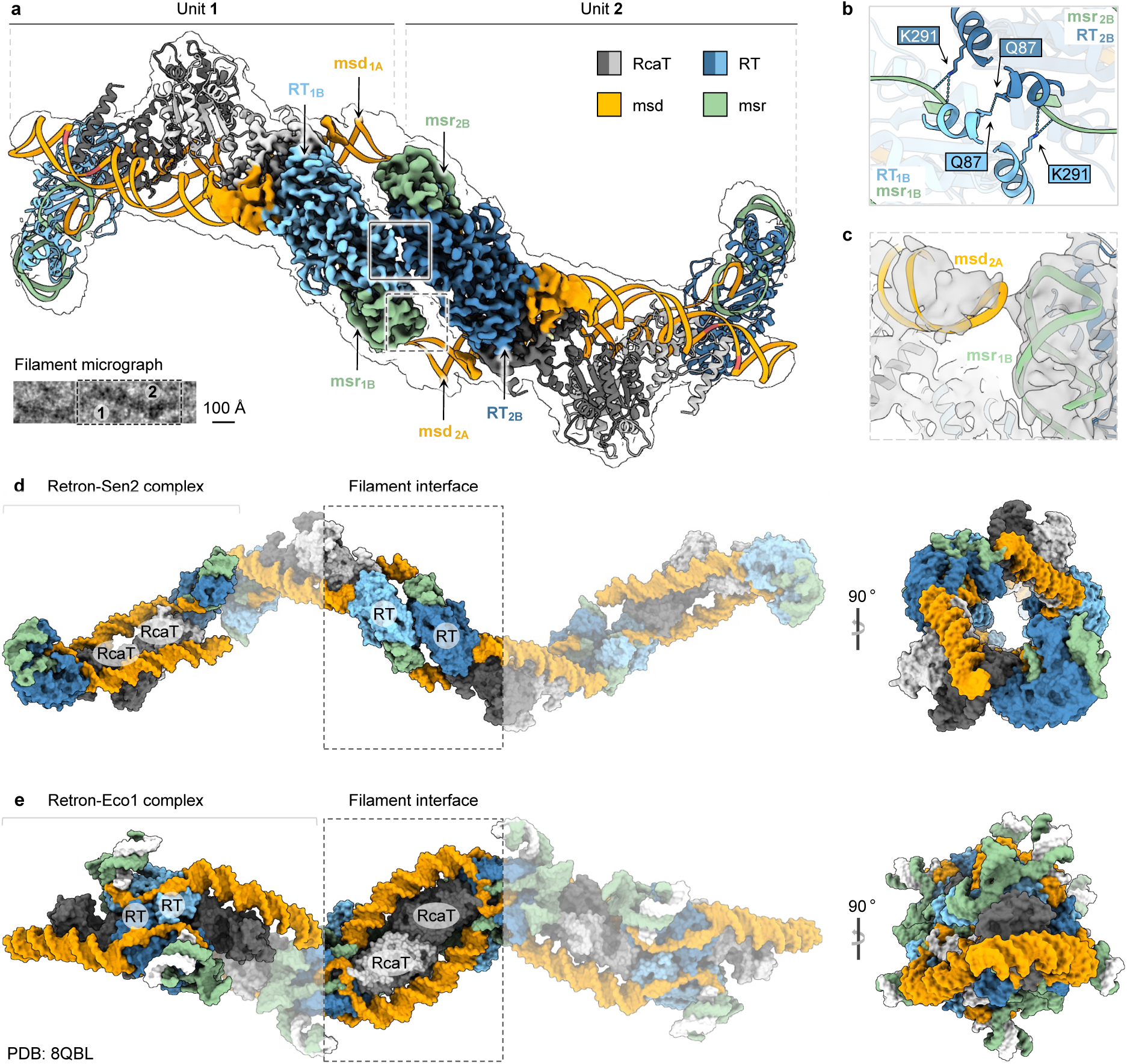
Retron-Sen2 and Retron-Eco1 filaments have inverted architectures. **a**, Model of a two-unit Retron-Sen2 filament fitted into the two-unit cryo-EM map (transparent outline; see Methods and Supplementary Data 6). At the centre of the filament, the locally refined filament interface map is coloured and its components labelled. The two repeating units are labelled 1 and 2 in both the 3D model and the representative micrograph (bottom left; scale bar, 100 Å). Boxes highlight the contacts detailed in panels b and c. Colours are consistent throughout all panels. **b**, msr appears to contact RT K291 at the filament interface, where a putative intermolecular RT Q87–Q87 contact is also observed. **c**, Interaction between msr from one repeating unit and msd from the adjacent unit may further stabilise the filament, although the local map quality does not permit detailed characterisation. Together, these contacts suggest a role for msr in filament assembly. **d**, Side and end-on views of a four-unit Retron-Sen2 filament model in surface representation (see Methods and Supplementary Data 6). The model approximates the architecture observed in the micrograph in panel a. The bracket marks the core toxin–antitoxin complex centred on an RcaT dimer (labelled RcaT–RcaT); the box highlights the filament interface centred on an RT-msDNA antitoxin dimer (labelled RT–RT). **e**, Corresponding views of a four-unit Retron-Eco1 filament (adapted from PDB 8QBL). In contrast to Retron-Sen2, the core complex centres on an RT-msDNA dimer, whereas the filament interface centres on an RcaT dimer. Components are colour-coded to match Retron-Sen2 for direct comparison, highlighting the inverted architectures of the two systems.

**Extended Data Figure 5.**
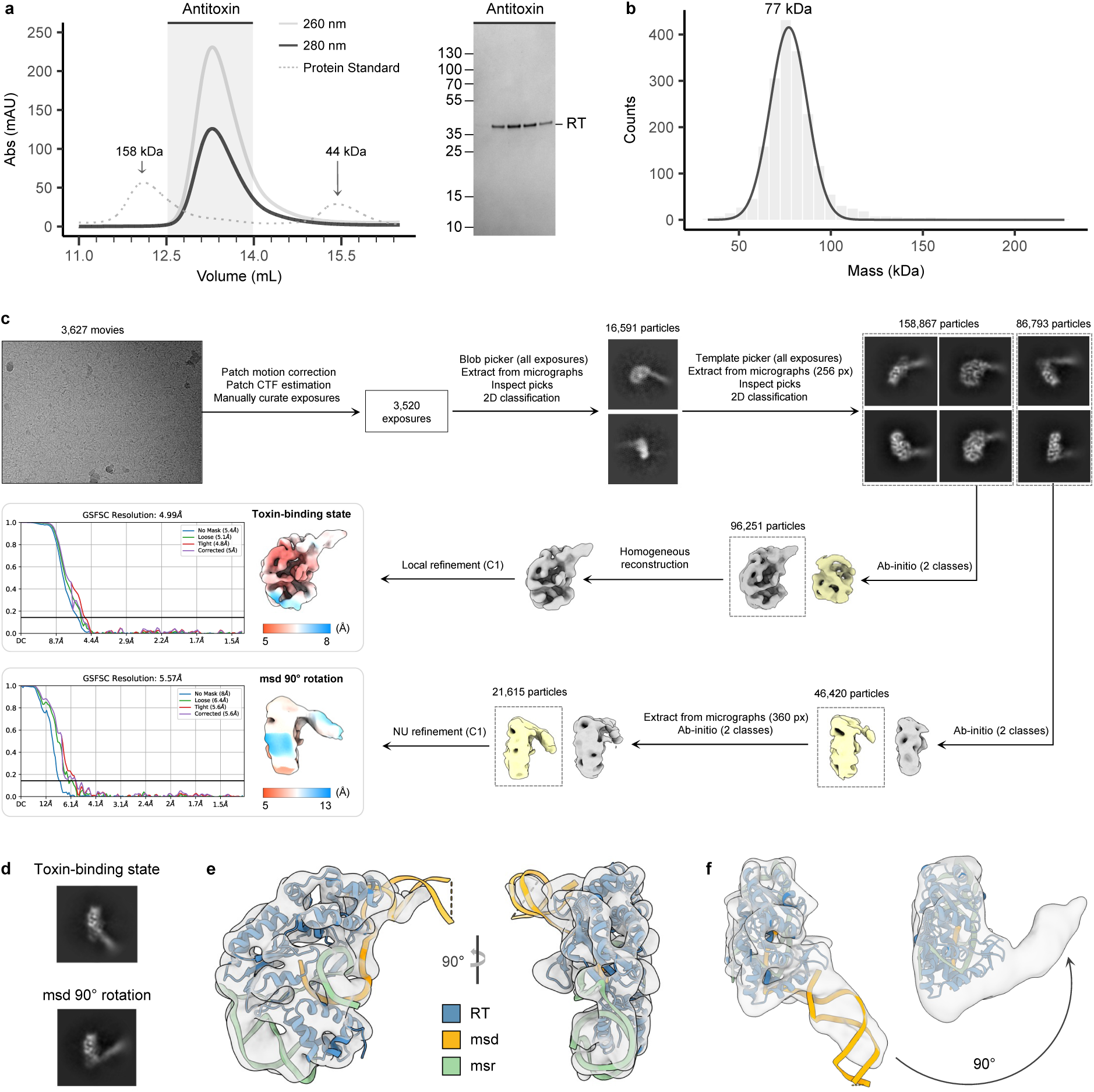
RcaT-free antitoxin adopts two distinct monomeric conformations. **a,** The SEC profile of purified antitoxin indicates that it is monomeric. SDS–PAGE of the corresponding peak fractions is shown at right. Light grey, dark grey and dotted traces indicate UV absorbance at 260 nm, 280 nm and protein standards, respectively. **b**, Mass photometry indicates a molecular mass of ∼77 kDa, consistent with a 1:1:1 complex comprising RT, msd and truncated msr. **c**, The Retron-Sen2 antitoxin was reconstructed in two monomeric conformations differing in msd orientation. The “toxin-binding state” was reconstructed to an estimated resolution of 4.99 Å, and the “msd 90° rotation” state to an estimated resolution of 5.57 Å. GSFSC curves and local resolution maps are shown for both reconstructions. **d**, 2D class averages of the two conformations in similar orientations illustrate the difference in msd orientation. **e**, The cryo-EM map of the toxin-binding state is shown with the fitted atomic model. In this conformation, msd adopts an orientation similar to that observed in the toxin–antitoxin complex. **f**, Cryo-EM maps of the toxin-binding state (left) and msd 90° rotation state (right) are fitted with their corresponding models. In the msd 90° rotation map, msd was omitted from the model, leaving residual density that reveals its orientation. The two states are positioned side by side to highlight the ∼90° rotation of msd between them.

**Extended Data Figure 6.**
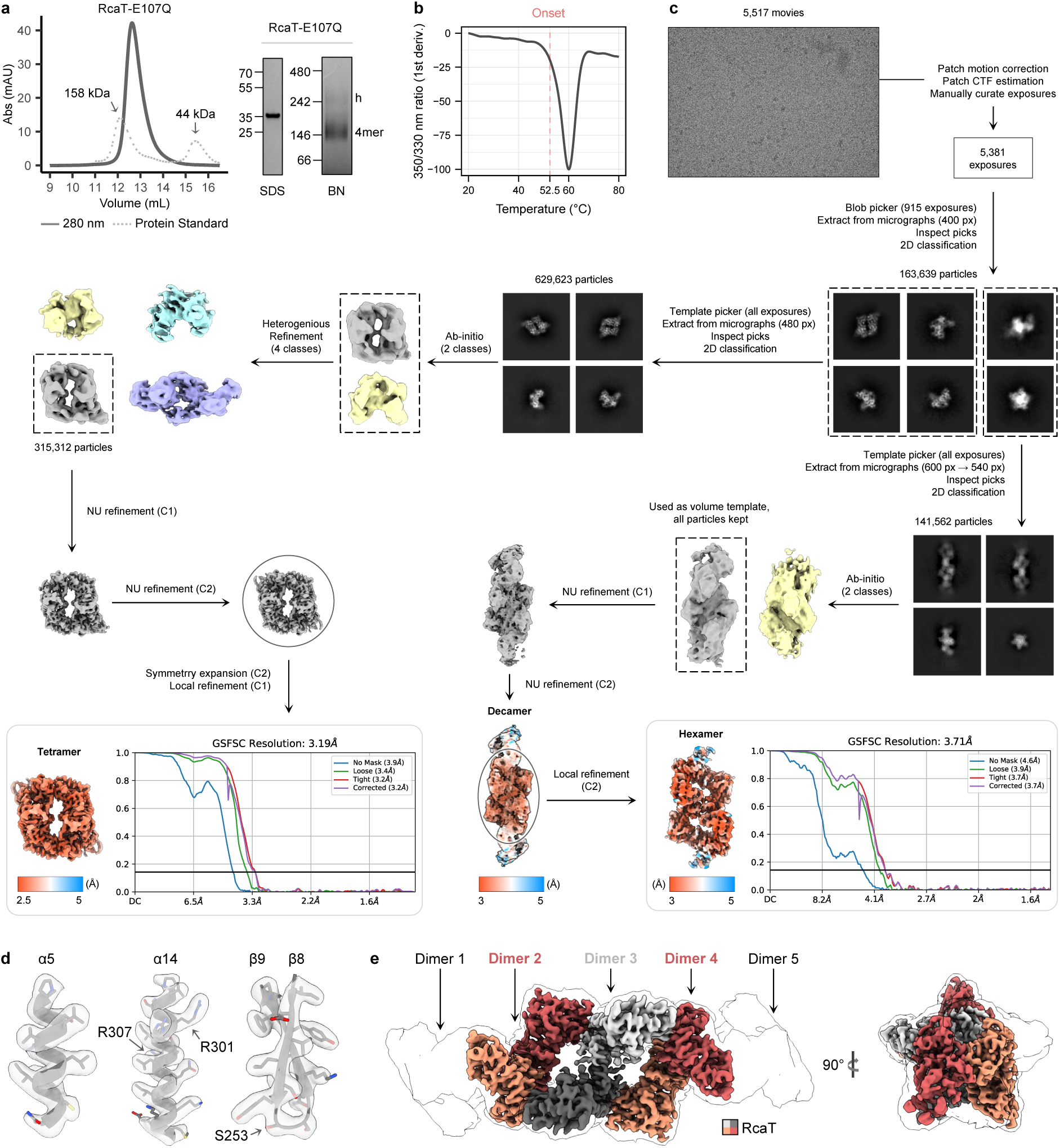
Structural characterisation of RcaT-Sen2. **a,** SEC shows that RcaT-Sen2 elutes predominantly as a tetramer. Solid and dotted traces indicate UV absorbance at 280 nm and protein standards, respectively. SDS–PAGE confirms protein purity, while blue-native PAGE (BN–PAGE) reveals higher-order oligomers (h) in addition to a dominant tetramer band (4mer). **b**, NanoDSF shows that RcaT-Sen2 begins to aggregate only above 52.5 °C, suggesting that its reported loss of toxicity at 37 °C in its native host^6^ does not result from thermal destabilisation. **c**, Cryo-EM data processing. RcaT-Sen2 adopts multiple oligomeric states. The tetramer was reconstructed to an estimated resolution of 3.19 Å. A decamer was also reconstructed, with local refinement of a hexamer yielding a map with an estimated resolution of 3.71 Å. GSFSC curves and local resolution maps are shown for both refined reconstructions. **d**, Close-ups show selected regions of the tetramer cryo-EM map (transparent surface) with the fitted atomic model. These regions mediate RcaT dimer–dimer assembly and include the β8– β9 hairpin and helices α5 and α14. Key residues are highlighted. **e**, The locally refined hexamer map (coloured) is shown within the decamer map (transparent). The top view reveals the characteristic five-pointed star shape of the decamer. The five stacked dimers forming the decamer are indicated.

**Extended Data Figure 7.**
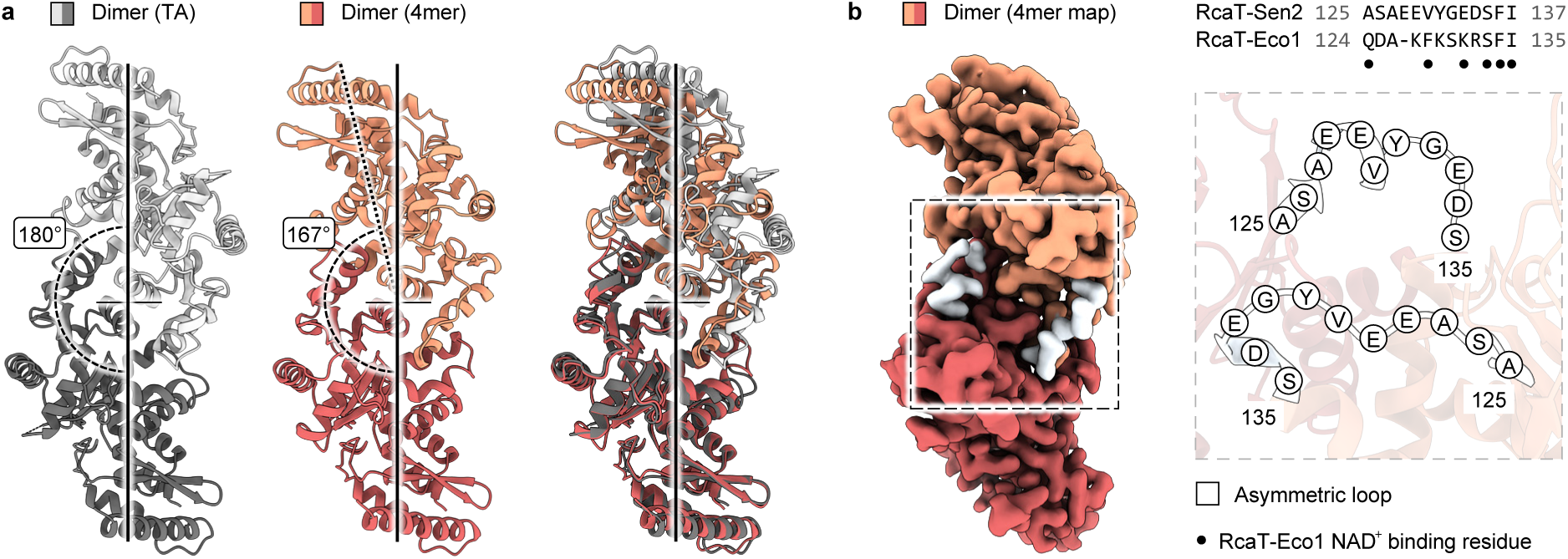
RcaT dimer–dimer assembly promotes a conformational change at the putative NAD⁺-binding site. **a,** The RcaT dimer observed in the toxin–antitoxin complex (light/dark grey) exhibits twofold symmetry, with the two subunits related by a 180° rotation. In contrast, the RcaT-Sen2 dimer observed in the tetramer (light/dark salmon) deviates from this symmetry by ∼13°. Superposition of the two dimers highlights this difference: the lower RcaT subunit aligns well (RMSD = 0.9 Å), whereas the upper deviates substantially. **b**, The asymmetry of the dimer within the tetramer originates from a loop spanning residues 125–137 (highlighted in white) near the hydrophobic core and active site of RcaT. A sequence alignment (generated using MAFFT) shows that several residues in the corresponding segment of RcaT-Eco1 contribute to NAD⁺ binding^13^, as indicated by black circles.

**Extended Data Figure 8.**
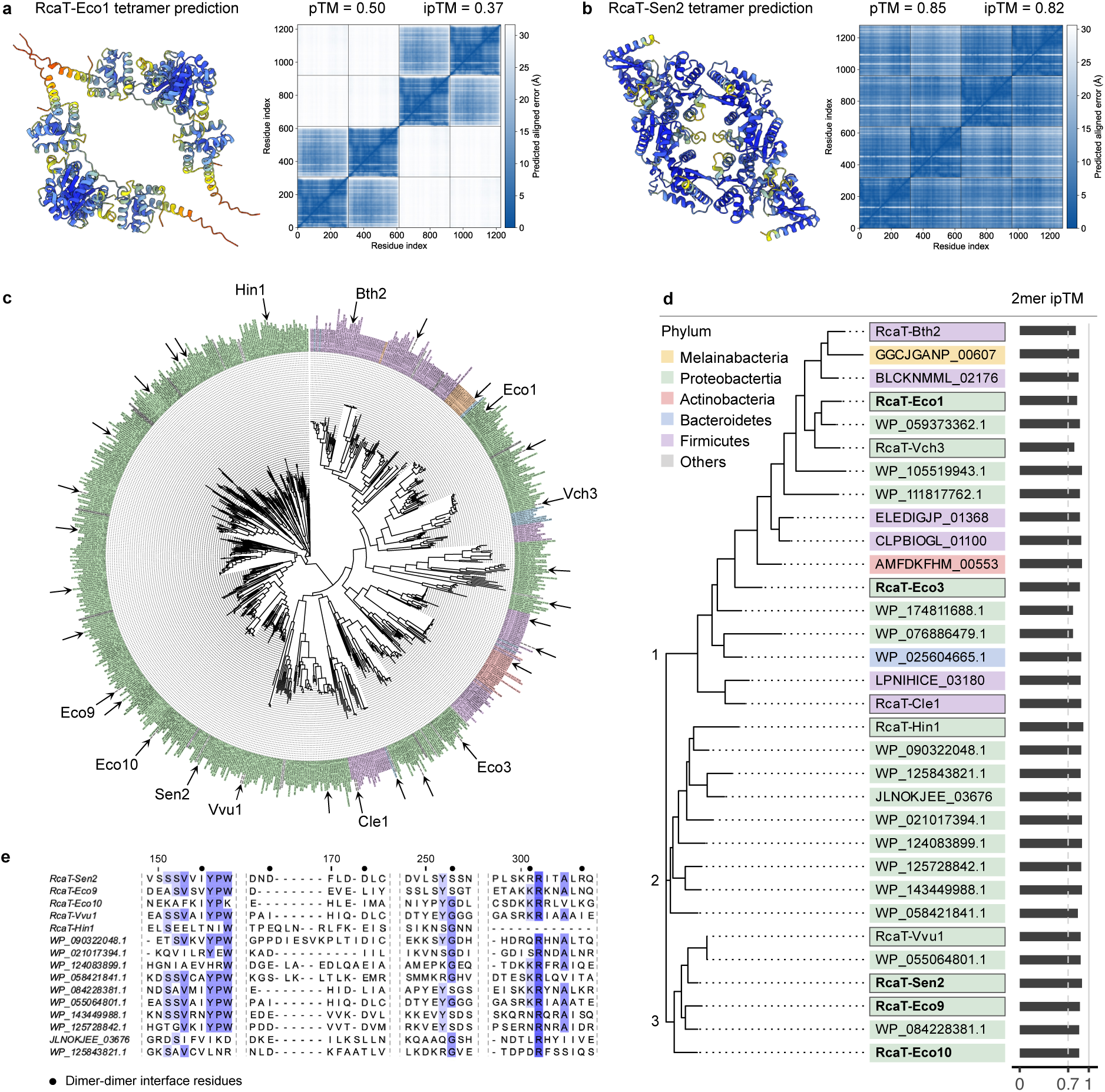
All RcaTs are predicted to form dimers, but only some form tetramers. **a,** AlphaFold fails to predict a confident homotetramer for RcaT-Eco1. The predicted model (left) consists of two separate dimers. The PAE plot and pTM and ipTM scores (right) indicate low confidence in the tetrameric arrangement. **b,** In contrast, AlphaFold confidently predicts a homotetramer for RcaT-Sen2. The predicted model (left) closely matches the experimental structure, and the PAE plot and pTM and ipTM scores (right) support the tetrameric arrangement. The examples in a and b are representative of unsuccessful and successful tetramer predictions, respectively, across the 32 RcaTs selected from the phylogeny. All predictions are provided in Supplementary Data 2-4. **c**, The 32 RcaTs selected for structural prediction from the 776-member RcaT phylogeny^6^ are marked by arrows and are representative of the tree. **d,** AlphaFold predicts that all selected RcaT homologues form dimers. Bars indicate ipTM scores for the predicted homodimers, with all predictions reaching high confidence (ipTM ≥ 0.7). Bacterial phyla are colour-coded as indicated in the inset and consistently applied in panels c and d. RcaTs tested for toxicity in Fig. 4c are outlined, and RcaTs with known toxicity profiles^6^ are highlighted in bold. **e,** Multiple sequence alignment of the 15 RcaTs predicted to form tetramers (performed using MAFFT and visualised with Jalview^56^). Black circles indicate residues involved in the dimer–dimer interface of RcaT-Sen2, the reference sequence. Overall, residues at the dimer–dimer interface are not conserved.

**Extended Data Figure 9.**
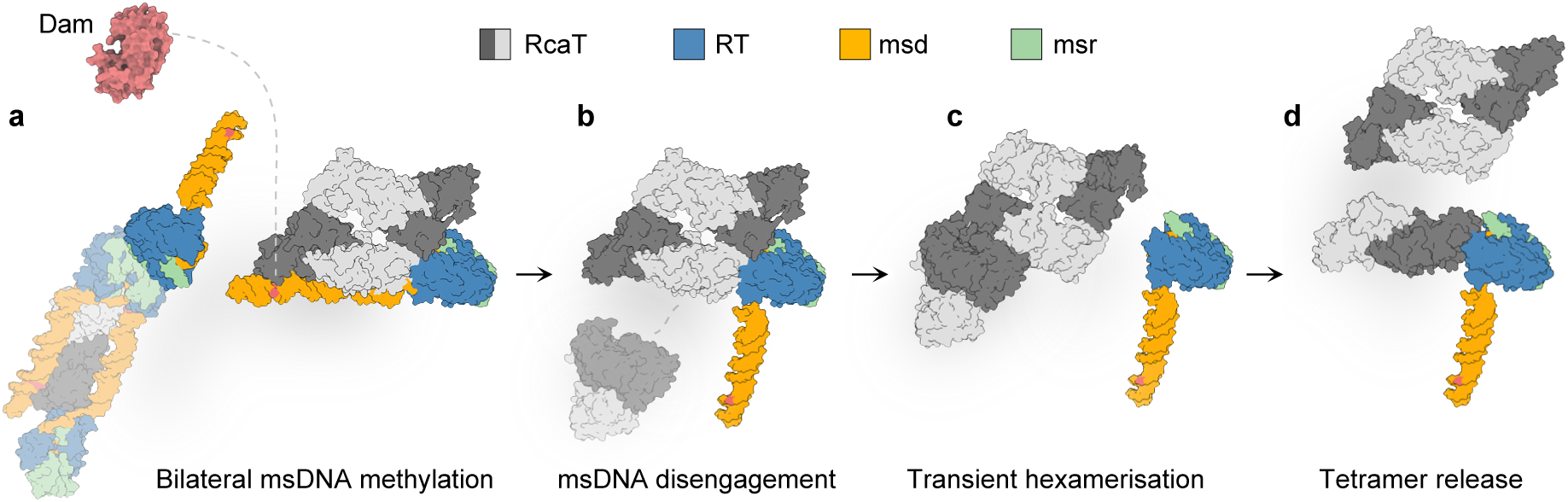
Putative model for methylation-mediated RcaT release. **a,** The model begins from the state shown in Fig. 4e: one msDNA has been methylated, allowing RcaT dimer–dimer assembly and displacement of one RT due to competition for helix α14. Dam methylates both msDNA molecules on either side of the RcaT dimer. **b**, Methylation of both msDNA molecules would promote their dissociation from both sides of the central RcaT dimer, creating space for a third incoming RcaT dimer. **c**, This third dimer could stack onto the central dimer, displacing the second RT and forming a transient hexameric assembly. **d**, The hexamer may subsequently dissociate into a released tetramer and a dimer that rebinds methylated antitoxin, consistent with the SEC of Fig. 2e. Incoming RcaT dimers are shown without their antitoxins for simplicity. Colour coding is indicated in the inset.

## Extended Data Tables

**Extended Data Table 1.** Cryo-EM data collection, refinement and validation statistics.

|  | #1 Retron-Sen2<br>toxin-antitoxin<br>complex<br>(EMD-100081)<br>(PDB 34PA) | #2 Retron-Sen2<br>2-unit filament<br>(EMD-100080) | #3 Retron-Sen2<br>filament<br>interface<br>(EMD-100090)<br>(PDB 34PL) | #4 Retron-Sen2<br>antitoxin<br>(EMD-100088)<br>(PDB 34PF) | #5 Retron-Sen2<br>antitoxin<br>alternative<br>conformation<br>(EMD-100079) | #6 RcaT-Sen2<br>tetramer<br>(EMD-100082)<br>(PDB 34PB) | #7 RcaT-Sen2<br>hexamer<br>(EMD-100083)<br>(PDB 34PC) | #8 RcaT-Sen2<br>decamer<br>(EMD-100084) |
| --- | --- | --- | --- | --- | --- | --- | --- | --- |
| **Data collection and processing** |  |  |  |  |  |  |  |  |
| Magnification | 130,000 | 130,000 | 130,000 | 130,000 | 130,000 | 130,000 | 130,000 | 130,000 |
| Voltage (kV) | 300 | 300 | 300 | 300 | 300 | 300 | 300 | 300 |
| Electron exposure (e<sup>-</sup>/Å<sup>2</sup>) | 58 | 58 | 58 | 51.1 | 51.1 | 46.36 | 46.36 | 46.36 |
| Defocus range (μm) | 0.75 – 2.5 | 0.75 – 2.5 | 0.75 – 2.5 | 0.75 – 2.5 | 0.75 – 2.5 | 0.75 – 2.5 | 0.75 – 2.5 | 0.75 – 2.5 |
| Pixel size (Å) | 0.68 | 0.68 | 0.68 | 0.68 | 0.68 | 0.68 | 0.68 | 0.68 |
| Symmetry imposed | C2 | C2 | C2 | C1 | C1 | C2 | C2 | C2 |
| Initial particle images (no.) | 2,554,504 | 2,920,302 | 2,920,302 | 5,287,109 | 5,287,109 | 2,861,659 | 2,590,747 | 2,590,747 |
| Final particle images (no.) | 262,357 | 147,852 | 147,852 | 96,251 | 21,615 | 315,312 | 141,562 | 141,562 |
| Map resolution (Å) | 3.37 | 3.55 | 3.29 | 4.99 | 5.57 | 3.19 | 3.71 | 3.78 |
| FSC threshold | 0.143 | 0.143 | 0.143 | 0.143 | 0.143 | 0.143 | 0.143 | 0.143 |
| Map resolution range (Å) | 2.8 – 47.1 | 2.8 – 42.7 | 2.8 – 30.4 | 4.4 – 8.12 | 5.5 – 13.9 | 2.0 – 7.5 | 2.0 – 7.0 | 2.0 – 35.8 |
| **Refinement** |  |  |  |  |  |  |  |  |
| Initial model used (PDB code) | - | - | - | 34PL | - | - | - | - |
| Model resolution (Å) | 3.8 | - | 3.9 | 7.1 | - | 3.4 | 4.1 | - |
| FSC threshold | 0.5 | - | 0.5 | 0.5 | - | 0.5 | 0.5 | - |
| Map sharpening B factor (Å<sup>2</sup>) | -50 | - | -50 | 0 | - | -100 | -50 | - |
| CC (mask) | 0.72 | - | 0.66 | 0.72 | - | 0.81 | 0.73 | - |
| **Model composition** |  |  |  |  |  |  |  |  |
| Non-hydrogen atoms | 15,180 | - | 12,310 | 4,083 | - | 10,352 | 14,634 | - |
| Protein residues | 1,254 | - | 938 | 311 | - | 1,280 | 1,812 | - |
| Ligands | 0 | - | 0 | 0 | - | 0 | 0 | - |
| **B factors (Å<sup>2</sup>)** |  |  |  |  |  |  |  |  |
| Protein | 154.08 | - | 129.94 | 387.54 | - | 132.21 | 98.30 | - |
| Ligand | - | - | - | - | - | - | - | - |
| **R.m.s. deviations** |  |  |  |  |  |  |  |  |
| Bond lengths (Å) | 0.003 | - | 0.004 | 0.003 | - | 0.002 | 0.002 | - |
| Bond angles (°) | 0.606 | - | 0.879 | 0.628 | - | 0.536 | 0.608 | - |
| **Validation** |  |  |  |  |  |  |  |  |
| MolProbity score | 1.14 | - | 1.41 | 1.89 | - | 0.89 | 1.31 | - |
| Clashscore | 3.39 | - | 5.38 | 12.24 | - | 1.48 | 5.36 | - |
| Poor rotamers (%) | 1.03 | - | 0.80 | 0 | - | 0.42 | 1.07 | - |
| **Ramachandran plot** |  |  |  |  |  |  |  |  |
| Favored (%) | 97.99 | - | 97.41 | 95.79 | - | 98.35 | 98.30 | - |
| Allowed (%) | 2.01 | - | 2.59 | 4.21 | - | 1.65 | 1.70 | - |
| Disallowed (%) | 0 | - | 0 | 0 | - | 0 | 0 | - |

